# Disrupting a Convergent Acetylation Circuit Collapses Leukemic Identity Across AML Subtypes

**DOI:** 10.64898/2026.07.13.737885

**Authors:** Anagha Deshpande, Cho-Ying Chiang, Marlenne Perales, Neha Niranjan, Neelam Sinha, Darren Finlay, Alexandra Stevens, Emily Zahn, Benjamin A. Garcia, Irmela Jeremias, Mark Wunderlich, Kristen Jensen-Pergakes, Akshata Udyavar, Amy Carr, Andrew R. Nager, Yanling Yang, Rabi Murad, Courtney Jones, Shawn O’Connell, Thomas Paul, Kristiina Vuori, Aniruddha J. Deshpande

## Abstract

Transcriptional condensates anchored by chromatin readers are increasingly recognized as organizing hubs for gene expression, but how their assembly and stability are regulated remains poorly understood. Here, we identify an acetylation-dependent feed-forward circuit that controls the integrity of the Super Elongation Complex (SEC), a key driver of transcriptional elongation. We show that the SAGA histone acetyltransferase catalytic subunits KAT2A/KAT2B license acetylation of both histone H3 lysine 9 (H3K9ac) and SEC components themselves, including ENL, AFF1, and AFF3. Loss of this dual acetylation activity, achieved via a cereblon-recruiting PROTAC (GSK983/GSK699), displaces the chromatin reader ENL from target loci, dissolves ENL-anchored transcriptional condensates, and disrupts SEC-dependent transcriptional output - linking histone and non-histone acetylation to the physical integrity of a core transcriptional machine.

Using genome-scale dependency data, we show that the SAGA complex is a selective chromatin dependency in acute myeloid leukemia (AML) AML and hematological malignancies and disrupting this feed-forward transcriptional circuit in AML demonstrates subtype independent antileukemia effects. KAT2A/B degradation drives potent, broad-spectrum antileukemic activity across genetically diverse AML cell lines, primary patient samples, and an isogenic KMT2A-rearranged model bearing cooperating oncogenic mutations, with H3K9ac loss concentrated asymmetrically at core AML oncogene loci such as MYC, MYB, and the HOXA cluster.

Together, these findings define an acetylation-dependent circuit governing SEC integrity and establish KAT2A/B degradation as a mechanism-based, pan-AML therapeutic strategy, with implications for transcriptional condensate regulation beyond leukemia.

**HIGHLIGHTS:**

- The SAGA complex is a selectively essential chromatin dependency across hematological malignancies and particularly in AML
- KAT2A/B degradation drives broad anti-leukemic activity across genetically diverse AML subtypes including chemo-refractory disease
- KAT2A/B degradation depletes H3K9ac at AML oncogene loci and dismantles ENL-anchored condensates
- KAT2A/B licenses regulation of super elongation complex acetylation and ENL interaction with SEC complex components

## INTRODUCTION

Acute myeloid leukemia (AML) is an aggressive hematological malignancy associated with a five-year survival of less than 30% ^1,2^. One of the key characteristics of AML is the clonal expansion of myeloid progenitors accompanied by a block in terminal differentiation, enhanced proliferative capacity, and resistance to apoptosis. In AML, the leukemic cells, and in particular the leukemic stem cell (LSC) population that drives disease maintenance and relapse ^3–5^, are defined not merely by their mutations that drive disease but by the gene expression program those mutations enforce. Many recurrent AML-associated gene mutations and gene fusions drive high-level, sustained transcription of a network of transcription factors and signaling modules that constitute leukemia cell identity and confer stem-like properties to AML cells ^6–11^ . There is now an increasing appreciation that the malignant transcriptional states that drive AML cell maintenance are not a passive consequence of AML-causing mutations, but instead reflect an actively maintained, chromatin-encoded program. Since AML cells can be profoundly and selectively dependent on these epigenetic programs, these dependencies have been termed transcriptional addiction ^12^ and the chromatin modulators involved in these programs are considered important therapeutic targets. Hence, the rationale is that disruption of epigenetic regulators that maintain leukemia-associated transcriptional programs can abrogate oncogenic transcription, exploiting the very feature that defines AML cell identity as its therapeutic vulnerability.

Based on this concept, several genetic studies - including ours - identified components of the SAGA (Spt-Ada-Gcn5 acetyltransferase) histone acetyltransferase (HAT) module – including the histone acetyltransferase KAT2A and the chromatin reader protein SGF29 as selective dependencies in AML ^13–16^. This convergence of multiple SAGA components on a shared AML dependency earmarks the SAGA complex as a therapeutic target and we hypothesized that targeted degradation of key components of this complex could disrupt the AML cell program with the selectivity and efficacy required for therapeutic benefit. In this study we conduct a comprehensive preclinical and mechanistic characterization of KAT2A/B PROTAC activity in AML. Our study reveals that KAT2A/B degradation drives selective suppression of leukemogenic transcription, leading to growth inhibition, induction of myeloid differentiation and suppression of leukemia stem cell programs in a broad panel of human AML cell lines, patient samples from genetically distinct mutational AML subtypes and models of chemo refractory AML. Mechanistically, KAT2A/B degradation causes loss of H3K9 acetylation and displaces key components of the super-elongation complex including ENL from chromatin, collapsing the ENL-anchored transcriptional identity program that drives leukemogenesis. Together, these findings establish KAT2A/B degradation as a potent mechanism-driven vulnerability in AML, providing a strong rationale for its therapeutic advancement.

## RESULTS

### The SAGA complex is selectively essential in hematological cancers and can be targeted by KAT2A/B degradation

Emerging evidence supports the notion that targeting chromatin complexes involved in selective control of leukemogenic transcription can produce profound therapeutic effects ^17,18^. Therefore, we first sought to identify protein complexes that are selectively essential for the fitness of hematological cancers using the Cancer Dependency Map (DepMap; Broad Institute, https://depmap.org/portal/). This dataset comprises whole-genome CRISPR-Cas9 screens conducted in > 1,000 cancer cell lines of diverse adult and pediatric tumor types. First, we used 718 chromatin modifying complexes from the CORUM database and defined a median single-sample gene set enrichment analysis (ssGSEA) score per complex to assess the CRISPR dependencies in cell lines from hematological cancers or AML specifically compared to all other cancer types. From these analyses, the SAGA complex emerged as one of the most significantly and selectively essential complexes in hematological cancers and also in AML compared to non-AML cell lines (Figure 1A, S1A) and Table S1. Focusing specifically on the SAGA complex, we then assessed dependency scores of this complex alone separately across cancer cell line lineages. This demonstrated that lymphoid and myeloid cancer cell lines displayed the highest SAGA dependency of all assessed tumor types (Figure 1B), whereas the closely related ATAC complex did not display such a hematological cancer-selective dependency (Fig. S1B). Next, pairwise comparisons of SAGA complex dependency showed that AML, leukemia broadly, and all hematological malignancies, each when compared to other cancer types - showed significantly higher SAGA dependency (p values 7.1 × 10^−4^, 6 × 10^−5^, and <10^−5^ respectively; Figure 1C). These findings position the SAGA complex as a compelling epigenetic vulnerability in hematological cancers, and particularly in AML.

**Figure 1.**
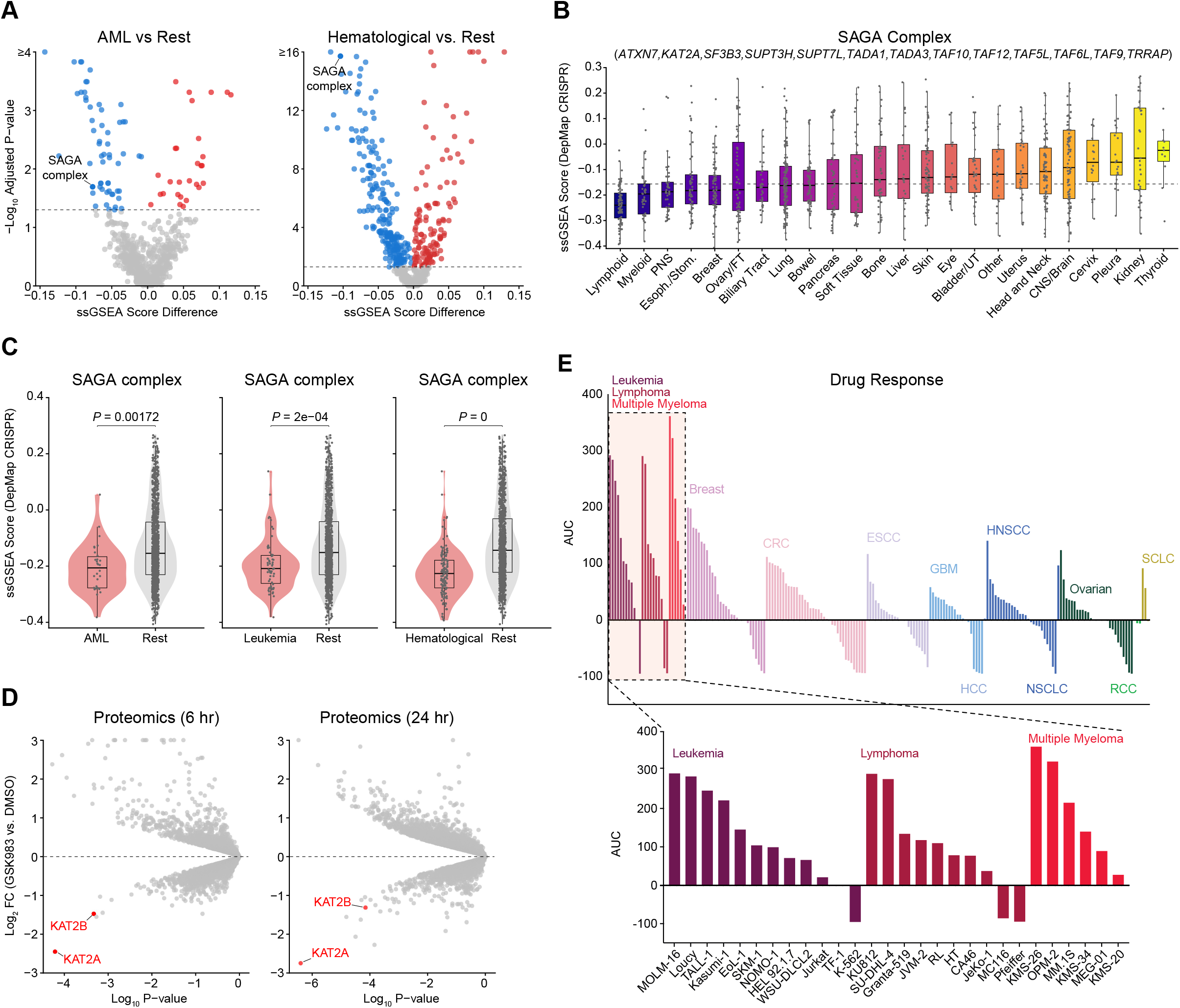
SAGA complex dependency is selective to hematological cancers and is targetable using a KAT2A/B degrader. **(A)** Volcano plot of ssGSEA scores for 718 CORUM protein complexes comparing AML vs. non-AML DepMap cell lines. The SAGA complex SAGA_476 from CORUM is labeled among the top AML-selective hits. Blue, AML-selective; red, selective in others; grey, NS. Dashed line: significance threshold. **(B)** Boxplot of SAGA complex CERES dependency scores (Avana SAGA Score) across cancer lineages. Lymphoid and myeloid lineages show the most negative scores, indicating the strongest dependency. Dotted line shows significance threshold. **(C)** Violin plots comparing SAGA_476 ssGSEA scores in AML vs. all others, leukemia vs. others, and hematological malignancies vs. other cancer types. Red: hematological cancer cell lines; grey: other cancer cell lines. Adjusted p-values are shown on top of each comparison **(D)** Volcano plots of global proteome changes in OCI-AML3 cells treated with GSK983 vs. DMSO at 6h (left) and 24h (right). Y axis represents log_2_ fold-change of GSK983 compared to DMSO whereas x axis represents log_10_ p-value. Each dot represents a protein; red: KAT2A and KAT2B; grey: all other proteins **(E)** Waterfall plots of GSK983 AUC-based sensitivity across ∼200 cancer cell lines colored by lineage. Upper panel: full panel of all the cell lines tested; lower panel: expanded view of the hematological cancer cell lines including leukemia, lymphoma and multiple myeloma. Y axis represents the Area Under Curve (AUC) whereas x axis represents the cell lines. CRC: Colorectal carcinoma; ESCC: Esophageal squamous cell carcinoma; GBM: Glioblastoma; HNSCC: Head and neck squamous cell carcinoma; HCC: Hepatocellular carcinoma; NSCLC: Non-small cell lung carcinoma; RCC: Renal cell carcinoma; SCLC: Small cell lung cancer

Targeted protein degradation has recently emerged as a powerful modality for inhibiting chromatin-associated oncogenic dependencies. The proteolysis-targeting chimera (PROTAC) GSK983 was developed as a degrader of KAT2A and KAT2B, the catalytic acetyltransferase subunits of the SAGA complex. To pharmacologically deplete KAT2A and KAT2B, we used a cereblon-recruiting proteolysis targeting chimera (PROTAC) comprising the selective PCAF/GCN5 bromodomain ligand GSK4027 conjugated to a thalidomide-based CRBN recruiter (termed GSK983 or its cis-(R,R) enantiomer GSK699^19^), hereafter referred to as the KAT2A/B PROTAC. Global proteomics in GSK983-treated OCI-AML3 AML cells at 6 and 24 hours demonstrated that KAT2A and KAT2B were among the most significantly and selectively depleted proteins at both timepoints, with minimal off-target proteome perturbation (Figure 1D and Table S2). We then profiled the activity of GSK983 in multiple cancer cell lines, including hematological, breast, colorectal, ovarian, lung, glioblastoma, head and neck, and esophageal cancer lines. Among ∼200 cancer cell lines spanning these diverse lineages, hematological malignancies including AML, ALL, lymphoma and multiple myeloma showed the greatest sensitivity to GSK983, with solid tumor lineages exhibiting substantially lower or near-complete insensitivity (Figure 1E and Table S3). These data provide direct evidence for the hematological cancer-selective dependency on the SAGA complex predicted by our DepMap analyses.

### KAT2A/B degradation impairs proliferation and clonogenicity across genetically diverse AML

Given that AML is a heterogeneous disease with distinct molecular groups and mutational subtypes, we sought to perform a detailed evaluation of GSK983 activity in cells from diverse AML subtypes. For this, we first tested the compound across a panel of human AML cell lines representing distinct driver oncogenes: MOLM-13 & NOMO-1 (KMT2A::MLLT3 positive), MV4-11 (KMT2A::MLLT4), OCI-AML3 (NPM1 mutant, DNMT3A mutant), and P31/Fujioka (PICALM::MLLT10 positive AML). GSK983 treatment led to a significant reduction in cell viability in a dose and time-dependent manner across all AML cell lines tested, with a progressive loss of cell growth observed from day 3 through 12 (Figure 2A) with a day 9 IC50 of 6-9 nM.

**Figure 2.**
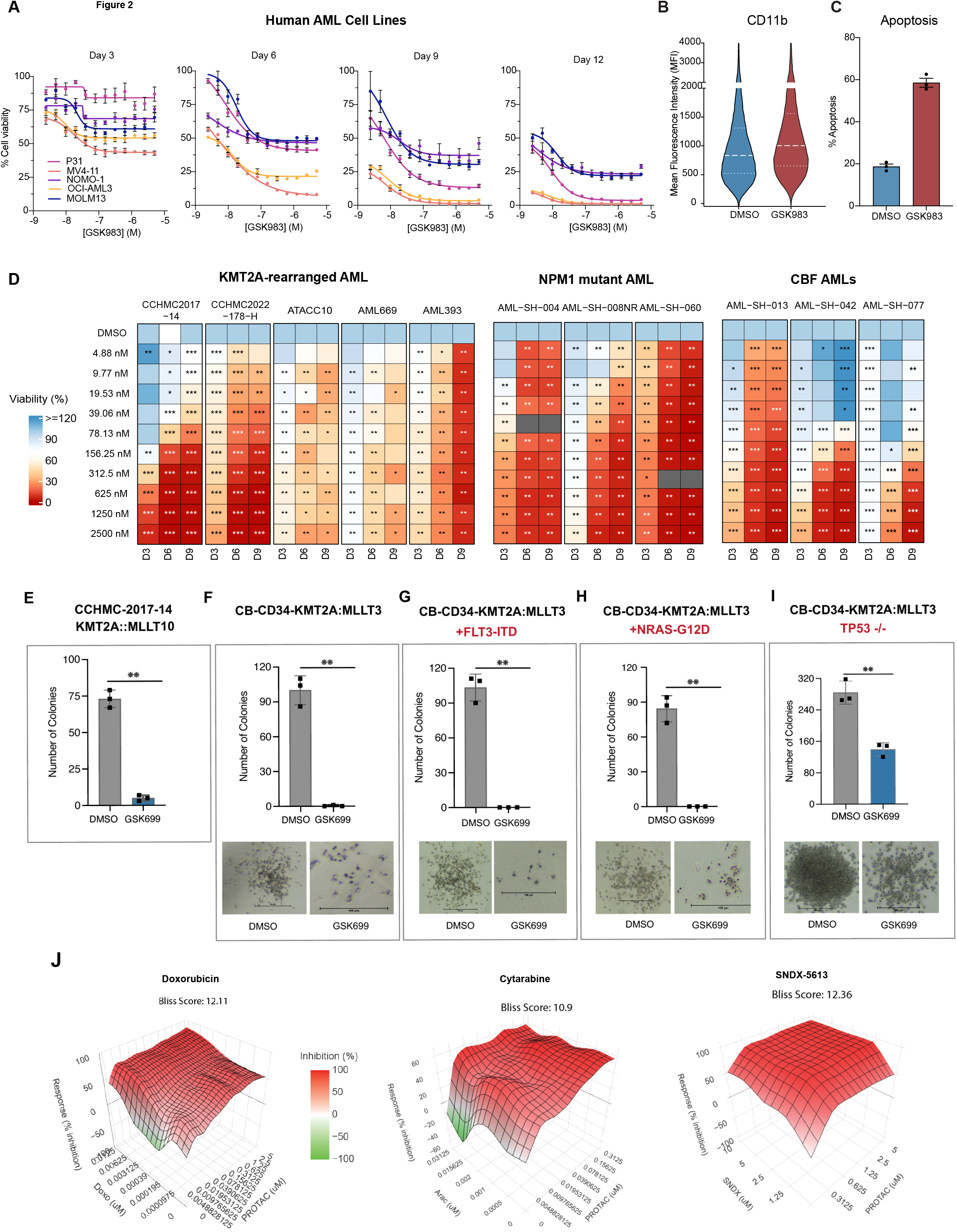
GSK983 impairs proliferation and clonogenic potential of genetically diverse AML models. **(A)** IC50 plots depicting dose response of 5 AML cell lines (P31, MV4-11, NOMO-1, OCI-AML3, MOLM-13) treated with GSK983 over the period of 12 days. Y axis shows percent cell viability normalized to DMSO control whereas x axis is Log Molar concentrations of GSK983. Left to right are the IC50 plot for Day 3, Day 6, Day 9 and Day 12. Data are mean ± SEM of n = 3 independent experiments. **(B)** CD11b surface marker expression (mean fluorescence intensity, MFI) changes in MOLM-13 treated with DMSO or 5 uM GSK983 for 72 hr. Violin plots show distribution across 3 independent replicates with Mean fluorescence intensity of APC representing CD11b expression on the y axis. Thin and thick dashed lines indicate quartiles and median respectively. **(C)** Apoptosis induction in MOLM-13treated with DMSO or 500 nM GSK983 for 7 days. Percent apoptosis quantified by flow cytometry is plotted on the y axis for DMSO and GSK983 treated cells. Data are mean ± SEM; n = 3. **(D)**Heatmap of GSK983 viability response across 11 primary AML patient-derived xenograft samples grouped by genetic subtype: KMT2A-rearranged (CCHMC2017-14, CCHMC2022-178H, ATACC 10, PDX-AML-669, PDX-AML-393), NPM1-mutant (AML-SH-004, AML-SH-008NR, AML-SH-060) primary AML samples, and core binding factor (CBF) AMLs (AML-SH-013, AML-SH-042, AML-SH-077). Samples were treated with GSK983 at concentrations from 4.88 to 2500 nM shown in rows and viability is shown at Days 3, 6, and 9 as columns. Color scale for percent viability normalized to DMSO is shown on the left of the heatmaps. Statistical significance versus DMSO calculated using the Mann-Whitney test is indicated as *p < 0.05, **p < 0.01, ***p < 0.001. **(E)** Colony-forming unit (CFU) assay for AML PDX sample. Number of CFUs per 3000 cells from CCHMC-2017-14 (KMT2A::MLLT10) cells treated with DMSO or 500 nM GSK699 for 10 days in H4100 semi solid media supplemented with 20% FBS and human cytokines are plotted on the y axis of the bar graph. Grey and blue bar represent colonies from DMSO and GSK699 treated cells respectively. N= 3 independent replicates; Error bars show mean ± SEM colony number; **p < 0.01 by unpaired t-test. **(F–I)** CFU assays for human cord blood derived CD34 cells transformed with KMT2A::MLLT3 alone (F), with FLT3-ITD (G), with NRAS-G12D (H), or with biallelic TP53 deletion (I) treated with DMSO or 500 nM GSK699 for 10 days. Colonies were counted on day 10 and total number of colonies per 2000 cells are plotted on the y axis of the bar graphs with grey bars representing DMSO and blue representing GSK699 treated colonies. N= 3 independent replicates; error bars show SEM values; **≤0.01. Below are the pictures of representative colonies from DMSO (left) and GSK699 (right) for each sample. Scale bar:100um. **(J)** Three-dimensional surface plots depicting drug combination responses between the KAT2A/B PROTAC (GSK983) and doxorubicin (left), cytarabine (center), or the menin inhibitor SNDX-5613 (right) in OCI-AML3 cells (n=3). Response is expressed as percent inhibition relative to DMSO control. Color scale indicates percent inhibition (red) or growth stimulation (green). Bliss synergy scores are shown above each plot.

GSK983 treatment also elicited a marked increase in CD11b expression relative to DMSO treated controls (Figure 2B and S2A), consistent with myeloid differentiation. In parallel, there was also a significant increase in Annexin V+ apoptotic cells in GSK983-treated AML cells (Figure 2C and S2B), indicating that growth suppression reflects a combination of differentiation and cell death.

We then extended our analysis to primary patient-derived AML samples. We profiled GSK983 on a total of 14 primary AML samples and patient-derived xenograft (PDX) specimens which included three major genetic subgroups: KMT2A-rearranged AML (n=4), NPM1-mutant AML (n=3), and core binding factor (CBF) AMLs (n=3) and others (n=3) (Table S4). Across all AML subgroups, GSK983 significantly suppressed AML cell growth compared to DMSO-treated controls with IC50 values ranging from sub-nanomolar to 310 nM at day 6, except for a patient sample with the PICALM::MLLT10 fusion that was insensitive (Figure 2D and Figure S2C). Notably, the antiproliferative activity of GSK983 was maintained across nearly all samples representing distinct cytogenetic lesions within each subgroup suggesting that it is not restricted to a specific AML mutation or fusion subtype.

Next, we evaluated the activity of GSK983 on clonogenicity of AML patient cells using the colony-forming unit (CFU) assay. GSK983 treatment of CCHMC-2017-14 AML cells from a patient with a highly refractory KMT2A::MLLT10 positive AML nearly eliminated their colony forming capacity (Figure 2E) - demonstrating the potent effects of GSK983 on AML cell clonogenicity. To investigate this further in a more defined genetic system, we used cord blood-derived CD34+ hematopoietic progenitors transformed with KMT2A::MLLT3 alone or in combination with cooperating oncogenic mutations (FLT3-ITD or NRAS-G12D) ^20^. Our results showed that GSK983 potently suppressed colony formation in all models, despite the additional oncogenic hits in this isogenic setting (Figures 2F–H). The striking suppression of colony forming units in the KMT2A::MLLT3 + FLT3-ITD and + NRAS-G12D contexts is particularly notable, as these mutations are associated with poor clinical outcomes and resistance to targeted therapies, suggesting that KAT2A/B dependence is maintained even in the setting of proliferative co-mutations. When we evaluated GSK983 activity in the context of an isogenic TP53 deleted setting – by biallelic TP53 deletion using CRISPR in KMT2A::MLLT3 transformed CD34 cells, the reduction in blast-like colonies was still significant (∼ 2-fold reduction) but not as dramatic as the FLT3-ITD and NRAS-G12D mutant settings (Figure 2I). Notably, these KAT2A/B degraders are only stable in culture for the first 2-3 days, yet the suppression of clonogenic growth in the 12-day CFU assay remained profound, indicating that even transient KAT2A/B degradation might be sufficient to irreversibly impair the leukemic and/or clonogenic capacity of AML progenitors.

Consistent with findings from the patient samples, GSK983 treatment also potently suppressed the formation of blast-like colonies in primary, secondary and tertiary replating from murine KMT2A::MLLT10 AML cells, with near-complete loss of blast-like colonies by week 3 (Figure S2D). Next, to determine the interaction of KAT2A/B degradation with agents currently used in the treatment of AML, we performed matrix drug combination assays in OCI-AML3 cells using GSK983 in combination with doxorubicin, cytarabine, or the menin inhibitor SNDX-5613. Of note, all three combinations yielded a strongly positive Bliss synergy score (doxorubicin: 12.11; cytarabine: 10.9; SNDX-5613: 12.36), indicating that KAT2A/B degradation may synergize with both standard-of-care chemotherapy and targeted epigenetic therapy in AML (Figure 2J). Together, these data establish that GSK983 exerts potent, broad-spectrum anti-leukemic activity and is particularly effective against leukemic progenitors.

### KAT2A/B degradation selectively reduces H3K9ac at AML oncogene loci and depletes stem-like cell populations in AML

To define histone modification landscape alterations mediated by the loss of KAT2A/B, we performed quantitative histone proteomics across 45 histone marks in OCI-AML3 cells treated with GSK983 or DMSO control. Of all modifications surveyed, our studies showed that H3K9ac was the only significantly altered mark, with a robust reduction following KAT2A/B degradation (Figure 3A and Table S5). This specificity for H3K9ac is consistent with this modification being the primary histone substrate of KAT2A/B, while also confirming that KAT2A/B degradation does not broadly disrupt the histone modification landscape, instead producing a highly targeted epigenomic perturbation.

**Figure 3.**
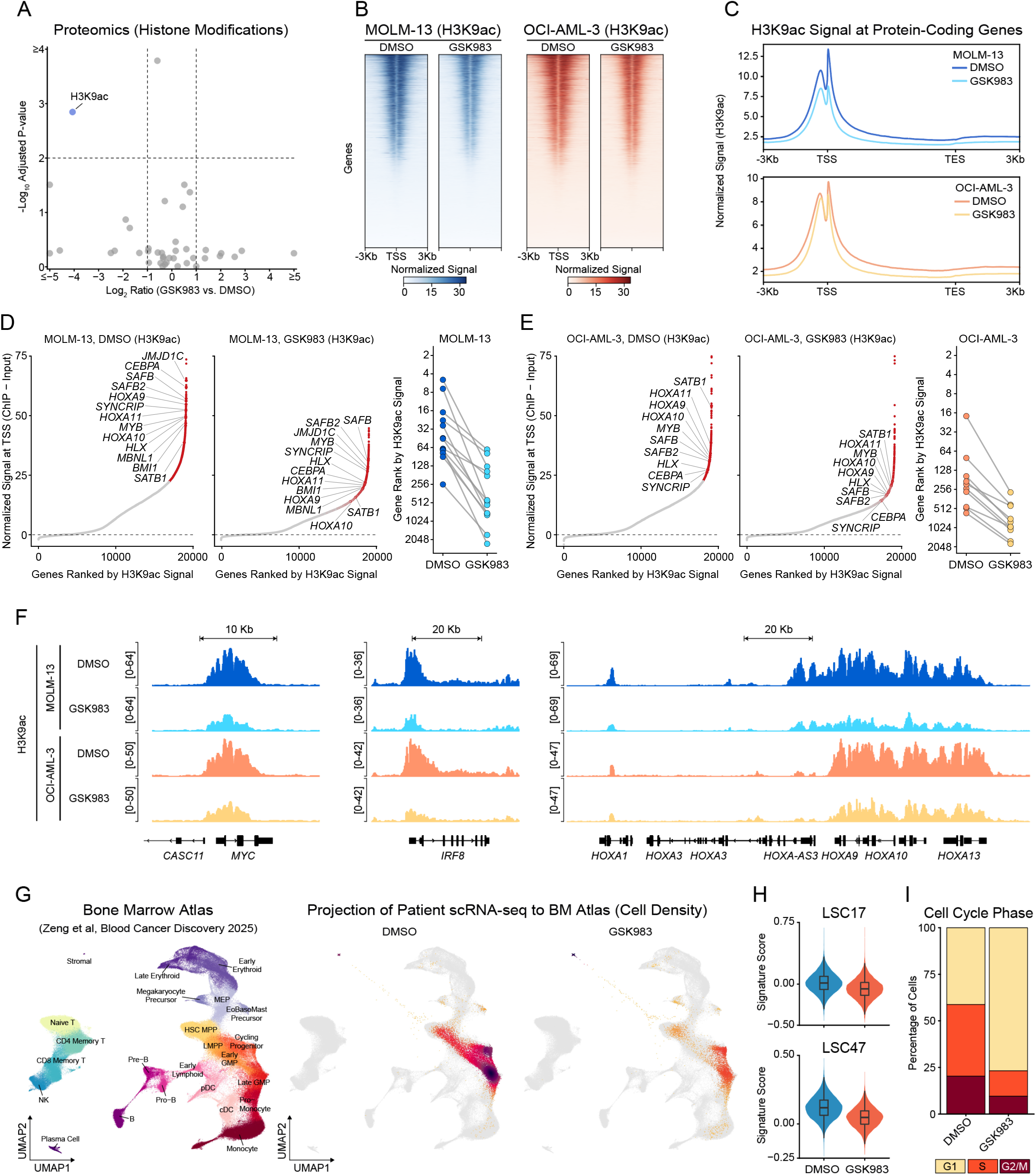
KAT2A/B degradation reduces H3K9ac at oncogene promoter loci and depletes primitive population in refractory AML. **(A)** Volcano plot depicting changes in histone post translational modifications using mass spectrometry. The log2 ratio of GSK983 treated vs DMSO (x axis) are plotted against −log10 adjusted p-value (y axis) for various histone modifications. Each dot represents a histone modification with H3K9ac in blue and unchanged modifications shown in grey. The data is obtained from 4 independent replicates. **(B)** Heatmaps of H3K9ac ChIP-seq signal (normalized) centered on transcription start sites (TSS ± 3 kb) of all protein-coding genes in MOLM-13 (blue) and OCI-AML3 (red) cells treated with DMSO or GSK983. Genes are ranked by decreasing H3K9ac signal in the DMSO condition. Color scale represents normalized ChIP-seq signal. **(C)** Meta-Gene analysis of H3K9ac ChIP-seq signal from −3 kb of TSS to +3 kb of transcription end site(TES) in MOLM-13 (top, blue) and OCI-AML3 (bottom, orange) cells treated with DMSO (dark) or GSK983 (light) are shown. Y-axis shows normalized H3K9ac signal. **(D-E)** Left and center: Rank-order plots of normalized H3K9ac signal at TSS for all protein-coding genes in MOLM-13 (D) and OCI-AML (E) cells treated with DMSO or GSK983. Key AML oncogenes are labeled and represented by red dots. Right: Connected slope graph showing the change in H3K9ac rank for the top-ranked AML oncogenes in DMSO following GSK983 treatment in MOLM-13 (blue) and OCI-AML3 (red). **(F)** Genome tracks on the integrative genomics viewer (IGV) of H3K9ac signal at the MYC, IRF8 or *HOXA* cluster in MOLM-13 (top, blue) and OCI-AML3 (bottom, orange) cells treated with DMSO (dark) or GSK983 (light) are shown. **(G)** scRNA-seq UMAP projections of AML sample treated with DMSO or GSK983. Left: normal human bone marrow reference atlas (Zeng et al., 2025). Middle: DMSO-treated PDX cells, ATACC 10 from a refractory relapsed AML patient, enriched for HSC, MPP, LMPP, and myeloid progenitor populations. Right: GSK983-treated ATACC 10 cells showing depletion of HSC/MPP/LMPP clusters and relative expansion of differentiated myeloid populations. **(H)** Violin plots showing LSC17 (top) and LSC47 (bottom) leukemic stem cell signature scores from scRNA-seq in ATACC 10 cells treated with DMSO in blue or GSK983 in red. Boxes indicate interquartile range; horizontal lines indicate median. **(I)** Stacked bar plots showing the percentage of cells in G1, S, and G2/M cell cycle phases in ATACC 10 cells treated with DMSO or GSK983 for scRNA-seq.

We performed chromatin immunoprecipitation followed by sequencing (ChIP-seq) in MOLM-13 and OCI-AML3 cells treated with GSK983 or DMSO to define genomic sites that had alterations in the H3K9ac chromatin mark. We observed that KAT2A/B degradation elicits a global reduction in H3K9ac signal across active gene promoters in both cell lines, with the most pronounced losses concentrated at the TSS-proximal regions of highly acetylated loci (Figures 3B and 3C). These data confirm that KAT2A/B degradation produces a genome-wide reduction in H3K9ac at promoter-proximal loci.

We then asked whether H3K9ac loss upon KAT2A/B degradation was uniform or preferentially affected specific gene programs in AML cells. Ranking all genes using the TSS-associated H3K9ac signal in AML cell lines (MOLM13 and OCI-AML3) revealed that promoters of core oncogenes that drive AML identity, oncogenesis and self-renewal - including *HOXA9, HOXA10, HOXA11, MYB, BMI1*, and *HLX* – had an asymmetrically high level of H3K9ac in the genome and GSK983 treatment most profoundly reduced the H3K9ac at the promoter-proximal sites of these oncogenic loci (Figure 3D-F). Specifically, the TSS of genes such as *HOXA9, HOXA10*, and *HOXA11* fell sharply in rank following GSK983 treatment, while other broadly active genes were comparatively less affected in both MOLM13 as well as OCI-AML3 cells (Figure 3D-F). This data suggests that KAT2A/B activity is disproportionately required to maintain the high acetylation state at these AML identity-defining promoter loci, and that their transcriptional output might therefore be particularly sensitive to KAT2A/B loss.

To determine whether H3K9ac loss at AML oncogene loci translates to functional changes in AML cell states, we performed single-cell RNA sequencing (scRNA-seq) on ATACC 10 AML cells, which are derived from a patient with highly chemorefractory, relapsed AML and harbor the KMT2A::MLLT4 fusion and NRAS G12A mutation – associated with dismal prognostic outcomes. Projection of the scRNA-seq transcriptomes onto a reference normal human bone marrow atlas ^21^ revealed that the ATACC 10 sample was enriched for cells resembling the transcriptomes of normal hematopoietic stem cell (HSC), multipotent progenitor (MPP), and lymphoid-primed multipotent progenitor (LMPP) populations as well as myeloid progenitors - a cellular composition reflecting the primitive, undifferentiated character of this highly refractory disease (Figure 3G, left and middle panels). Following GSK983 treatment, there were profound depletions of the HSC, MPP and LMPP-like populations accompanied by significant reduction in transcriptional signatures associated with leukemia stem cells (Figure 3G,right panel) and poor survival outcomes in AML (LSC17 ^22^) as well as in pediatric AML (LSC47 signature ^23^) (Figure 3H). GSK983-treated ATACC 10 AML cells also exhibited an increased G1 and concomitantly reduced S-phase, indicative of cell cycle arrest (Figure 3I). These findings demonstrate that KAT2A/B degradation can attenuate the leukemic stem cell transcriptional program in primary AML cells, eliminating primitive AML cell populations even in the context of refractory disease.

### KAT2A/B degradation suppresses MYC transcriptional output and leukemia stem cell programs in AML

Given the importance of KAT2A/B inhibition/degradation as a novel therapeutic modality, we sought to characterize the transcriptional consequences of GSK983 treatment in AML in detail using 5 AML cell lines (OCI-AML3, MOLM13, HL60, P31 Fujioka and MV4-11). Principal component analysis of bulk RNA-seq data from five AML cell lines spanning these diverse genetic backgrounds revealed that GSK983 treatment consistently separated cells from their DMSO counterparts along a common principal component (Figure 4A), suggesting a shared transcriptional response independent of the underlying oncogenic driver. Indeed, a detailed interrogation of gene overlap across all five cell lines identified a core of 859 genes commonly downregulated in all five AML cell lines (Figure 4B and Table S6). Among the common downregulated GSK983 core response genes there was a strong enrichment for oncogenic and myeloid transcription factors (*MYC, MYB, MYBL2, IRF8*), components of the ribosome biogenesis pathway (*NCL, NOP58, NOP56, DKC1*), enzymes involved in cholesterol (*HMGCR, HMGCS1, SQLE, FDFT1*) and fatty acid synthesis (*FASN, SCD, FADS1/2, ACACA*), as well as glycolytic enzymes (*LDHA, LDHB, ENO1, PGK1, PFKP*) (Fig. 4B). Inhibitors of chromatin-associated complexes such as DOT1L and the Menin potently downregulate leukemogenic homeobox genes, a mechanism relevant to HOX-overexpressing AML such as KMT2A-rearranged or nucleophosmin 1 (NPM1) mutant AML. Similarly, GSK983 treatment caused a significant downregulation of HOXA cluster oncogenes such HOXA9, HOXA10 and HOXA13 in AML cells driven by high HOXA gene expression (OCI-AML3, MOLM13, P31-Fujioka and MV4-11) (Fig. 4B-C and Table S6), but in contrast to DOT1L or Menin inhibitors, KAT2 degradation showed highly potent antiproliferative activity even in AML that do not display HOX gene expression, such as the core binding factor (CBF) AMLs. A transcriptional response common to all AML cell lines examined, regardless of genetic subtype, was the downregulation of MYC as well as MYC target genes together with a concomitant upregulation of myeloid differentiation markers including ITGAM and CD86 as well as interferon-stimulated genes (ISGs). This shared transcriptional response identifies MYC program suppression as a subtype-agnostic consequence of KAT2A/B degradation that may underlie its broad anti-leukemic activity.

**Figure 4.**
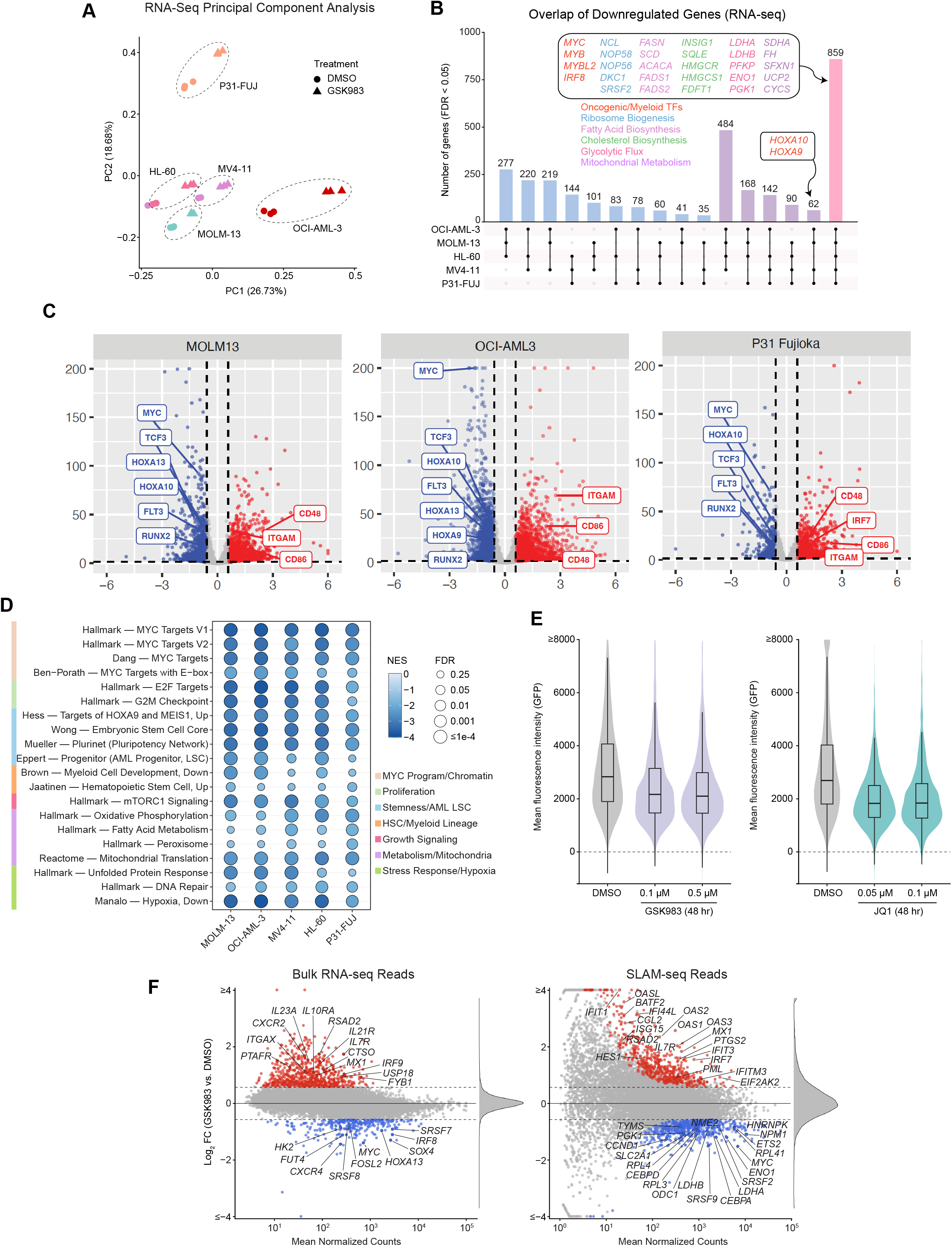
KAT2A/B degradation downregulates the MYC transcriptional program and oncogenic stemness signatures across genetically diverse AML. **(A)** Principal component analysis (PCA) of bulk RNA-seq data from five AML cell lines (OCI-AML3, MOLM-13, MV4-11, HL-60, P31-FUJi) treated with DMSO (circles) or GSK983 (triangles). Each cell line is shown in a distinct color with 95% confidence ellipses. PC1 (26.73%) and PC2 (18.68%) are shown. **(B)** UpSet plot showing the overlap of significantly downregulated genes (FDR < 0.05) across five AML cell lines treated with GSK983 for 48hr. Bar height indicates the number of genes in each intersection set. Colored gene names in the box denote their pathway mentioned with the matching color under the box. **(C)** Volcano plots showing bulk RNA-seq from MOLM-13 (left) OCI-AML3 (middle) and P31-Fujioka (right) cells treated with GSK983 or DMSO for 48hr. X-axis represents log_2_ fold-change compared to DMSO and the y-axis shows −log_10_ adjusted p-value. Significantly downregulated genes are marked with blue, significantly upregulated genes with red and non-significant ones are in grey. Dashed lines indicate significance thresholds. **(D)** GSEA bubble plot showing normalized enrichment scores (NES) for curated gene sets across five AML cell lines across the columns. Size of the circles represents FDR whereas color represents NES on a scale from 0 to −4 . Color stick on the left of plot indicates gene sets grouped by their biological pathways. Color for each group are shown in the legend on the right **(E)** Violin/box plots of mean GFP fluorescence intensity (MFI) from a MYC-GFP transcriptional reporter cells from *in vitro* transformed mouse leukemia harboring PICALM::MLLT10 fusion treated for 48 hours with increasing doses of GSK983 (left: 0.1 µM and 0.5 µM) or JQ1 (right: 0.05 µM and 0.1 µM), compared to DMSO control. Dashed line indicates zero fluorescence baseline whereas solid line inside box plot indicates median of MFI. N=3; grey: DMSO, Purple: GSK983 treated and teal: JQ1 treated. **(F)** MA plots of bulk RNA-seq (left) and SLAM-seq (right) in OCI-AML3 cells treated with 150 nM GSK983 or DMSO for 4 hr. Each point represents a gene; x-axis shows mean normalized counts (log_10_ scale) and y-axis shows log_2_ fold-change. Significantly upregulated genes are shown in red; downregulated genes in blue and unchanged in grey. The dotted lines indicate the cut-off value.

Specifically, gene set enrichment analysis (GSEA) of the bulk RNA-seq data across all five AML cell lines revealed a strikingly consistent pattern of pathway suppression. The most significantly and consistently depleted gene sets were MYC target signatures, E2F targets, and G2M checkpoint genes - indicating coordinate suppression of MYC-driven proliferative transcription (Figure 4D). Consistent with the role of the SAGA complex in AML LSC regulation, stemness and AML LSC programs were also uniformly depleted across all lines. Finally, suppression of metabolic pathways including oxidative phosphorylation, mTORC1 signaling, fatty acid metabolism, and mitochondrial translation was also uniform across all five AML cell lines (Figure 4D), consistent with a broad suppression of mitochondrial respiratory capacity. To assess whether KAT2A/B degradation disrupts cellular bioenergetics, we performed Seahorse XF Mito Stress Test assays in MOLM-13 and OCI-AML3 cells treated with GSK983 or DMSO for 4 hours. In these assays, we observed that GSK983-treated cells showed a marked reduction in basal and maximal mitochondrial oxygen consumption rate (OCR) as well as extracellular acidification rate (ECAR) relative to DMSO controls in both cell lines, indicating that acute KAT2A/B degradation broadly impairs oxidative phosphorylation and glycolytic capacity in AML cells. (Figure S3A-D).

To directly validate MYC transcriptional suppression at the protein level, we used AML cells derived from murine bone marrow in which the murine Myc gene is tagged with a carboxy-terminal green fluorescence protein (GFP), acting as a reporter of MYC expression. In these CALM-AF10 transformed Myc-GFP cells, GSK983 treatment produced a significant and dose-dependent reduction in the Myc-GFP signal as measured using flow cytometry, with marked suppression at 0.5 µM (Figure 4E, left).

Next, we sought to determine whether these transcriptional effects were a direct consequence of GSK983 treatment or simply reflected a differentiation state that resulted from KAT2A/B loss. For this, we performed thiol (<u>S</u>H)-linked <u>a</u>lkylation for the <u>m</u>etabolic <u>seq</u>uencing of RNA (SLAM seq) which enables discrimination of newly transcribed from pre-existing mRNA transcripts, allowing direct measurement of nascent transcription rather than steady-state mRNA levels. In contrast to the 48-hour drug exposure we used for the bulk-RNA seq experiments described above, here we treated OCI-AML3 cells with GSK983 for only 4 hours followed by 1 hour of RNA labeling. This nascent RNA sequencing experiment demonstrated a significant reduction in MYC transcripts as well as of mRNAs encoding core glycolytic enzymes (*LDHA, LDHB, ENO1, PGK1*) and ribosomal protein genes (*RPL3, RPL4, RPL41*) (Figure 4F). These data demonstrate that transcriptional modulation of key GSK983 target genes is an early and possibly direct consequence of KAT2A/B degradation, preceding any overt differentiation of AML cells. Notably, across all four orthogonal platforms - RNA-seq, SLAM-seq, ChIP-seq, and total proteomics - the interferon regulatory transcription factor *IRF8* emerged as one of the most consistently and significantly downregulated targets following KAT2A/B degradation, ranking above even *MYC* in several datasets, suggesting that KAT2A/B activity is required to sustain the aberrant expression of this myeloid identity factor at levels that support the leukemic progenitor state.

Taken together, this pan-AML convergence on common pathway suppression, spanning AML cell lines with distinct genetic drivers argues that KAT2A/B activity is a shared epigenomic dependency for the maintenance of leukemic stem cell identity, MYC-driven proliferation, and metabolic homeostasis in AML.

### KAT2A/B degradation disrupts ENL and RNA Poll II occupancy at asymmetrically acetylated AML oncogene promoters

Given that KAT2A/B degradation rewires the chromatin landscape at the promoters of AML oncogenes, we reasoned that genetic perturbation of epigenetic regulators might reveal downstream effectors required for GSK983 activity, offering insights into the mechanism of action of KAT2A/B degradation in AML. For this, we performed a focused CRISPR screen using our previously reported custom-made pooled library of sgRNAs targeting 644 epigenetic modulators ^15^ in the MOLM-13-Cas9 cell line. (see Figure 5A for schematic). Analysis of the CRISPR screen using Model-based Analysis of Genome-wide CRISPR-Cas9 Knockout (MAGeCK)^24^ identified several genes whose deletion conferred resistance to GSK983, i.e., the sgRNAs were positively selected in the drug treated compared to the vehicle-treated arm. Among the top genes whose loss conferred resistance to GSK983 - was KAT2A, indicative of on-target GSK983 activity. Strikingly, the chromatin reader protein *ENL* (MLLT1), together with *KMT5A, PHF23, UBE2A*, and *NSD1* were also among the top resistance hits (Figure 5B-C and Table S7). ENL is a chromatin reader protein that binds to acylated marks of actively transcribed genes on histone 3 - including H3K9ac – and recruits the super elongation complex (SEC). Since the reader activity of ENL has been shown to be important to sustain oncogenic transcription programs in AML ^25,26^, we wanted to ask whether KAT2A/B degradation and H3K9ac loss alter ENL chromatin occupancy. To address this, we performed CUT&RUN for ENL in GSK983-treated OCI-AML3 cells. Strikingly, gene promoters that lost H3K9ac upon GSK983 treatment showed coordinate diminution of ENL occupancy as well as total and serine 2 phosphorylated (Ser2P) Pol II (Figure 5D), with the corresponding genes downregulated at the transcript level (Figure 5E), indicating that H3K9ac loss is coupled to displacement of the ENL-tethered SEC complex and collapse of the RNA polymerase machinery at these loci.

**Figure 5.**
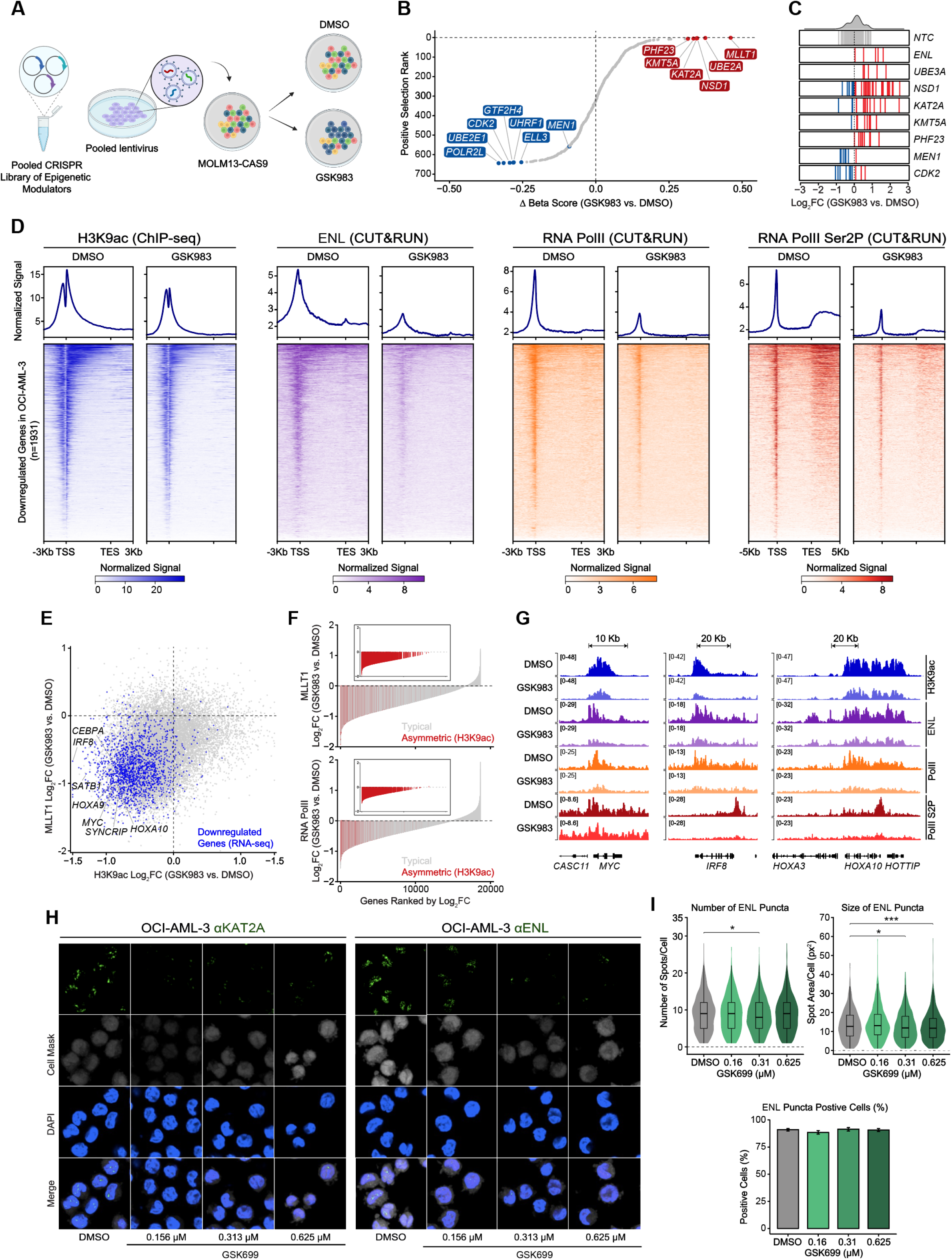
KAT2A/B degradation collapses ENL-anchored chromatin programs at oncogenic loci. **(A)** Schematic of the pooled CRISPR screen in the presence of GSK983: MOLM-13 Cas9-expressing cells were transduced with an epigenetic sgRNA library targeting ∼644 epigenetic and chromatin-associated genes, then split into DMSO and GSK983 treatment groups for 14 days. sgRNA abundance was quantified by next generation sequencing and differential enrichment was scored as Δ beta score. **(B)** Scatter plot of CRISPR screen results showing Positive Selection Rank (y-axis) versus Δ beta score (GSK983 vs. DMSO, x-axis). Resistance hits (positive Δ beta, red): *MLLT1, UBE2A, NSD1, KAT2A, KMT5A, PHF23*. Sensitizer hits (negative Δ beta, blue): *POLR2L, UBE2E1, CDK2, GTF2H4, UHRF1, ELL3, MEN1*. Dashed lines indicate Δ beta = 0. **(C)** Individual sgRNA log_2_FC (GSK983 vs. DMSO) depicted by waterfall distributions for selected screen hits and non-targeting controls (NTC). Each line represents one sgRNA with red and blue representing enriched and depleted respectively whereas grey representing unchanged NTC sgRNAs. Genes shown are NTC, *MLLT1, UBE2A, NSD1, KAT2A, KMT5A, PHF23, MEN1, CDK2*. **(D)** ChIP-seq and CUT&RUN heatmaps showing enrichment of H3K9ac (ChIP-seq), ENL, total RNA Pol II, and RNA Pol II Ser2P (all CUT&RUN) in OCI-AML3 cells treated with DMSO or GSK983, centered on the TSS of 1,931 GSK983-downregulated genes and ranked by DMSO signal. Regions span −3 kb to TSS to TES to +3 kb for H3K9ac, ENL, RNA Pol II and −5 kb to TSS to +5 kb of TES for RNA Pol II Ser2P. Metagene aggregate profiles are shown above heatmaps. Color scales indicate normalized signal. **(E)** Scatter plot demonstrating H3K9ac log_2_FC (x-axis) versus ENLlog_2_FC (y-axis) across the genome in OCI-AML3 cells treated with GSK983 when compared to DMSO control. DMSO. Each dot represents a gene with blue dots representing genes downregulated in RNA-seq. Select AML oncogenes are labeled. Dashed lines indicate log_2_FC = 0. **(F)** Waterfall plots showing ENL log_2_FC (top panel) and RNA Pol II log_2_FC (bottom panel) at genes ranked by their log_2_FC. Red and grey lines indicate genes with asymmetric loss and genes with typical H3K9ac changes respectively. Insets show expanded view of the asymmetric H3K9ac gene subset. **(G)** Integrative Genomics Viewer tracks (IGV) for H3K9ac (blue), ENL (purple), RNA Pol II (orange), and RNA Pol II Ser2P (red) in OCI-AML3 cells treated with DMSO (dark) or GSK983 685 (light) at three oncogenic loci: *CASC11*/*MYC* (left), *IRF8* (center), and *HOXA cluster* (right). y-axis 686 ranges of peaks are indicated per track; scale bars are shown above. **(H)** Immunofluorescence (high content imaging) indicating changes in condensate formation of 688 ENL after KAT2A degradation, GCN5/KAT2A (left, green) and ENL (right, green) in OCI-AML3 689 cells treated with DMSO, 0.156 µM, 0.313 µM, or 0.625 µM GSK699. Rows show results of 690 staining with antibody against KAT2A and ENL (GFP, top row), Cell Mask (grey, second row), 691 DAPI (blue, third row), and merged channels (bottom row). **(I) High-content immunofluorescence quantification of ENL nuclear condensates in OCI-AML3 693 cells treated with increasing doses of GSK699**. Percentage of ENL puncta-positive cells across 694 different drug conditions (DMSO, 0.16, 0.31, and 0.625 µM GSK699) are shown in the bottom 695 panel. *p < 0.05; ***p < 0.001 using the Wilcoxon rank sum test.

Of note, the loss of ENL and Pol II was most profound at gene promoters bearing asymmetrically high H3K9ac in AML cells including at key AML driver oncogene loci (Figure 5F-G).

The maintenance of transcription of lineage identity genes or oncogenic programs has been shown to be mediated through phase-separated transcriptional condensates which can compartmentalize and concentrate the transcription machinery at key genomic loci. Recent studies have shown that ENL forms transcriptional condensates by binding acetylated histones through its YEATS domain and recruiting the SEC to drive Pol II elongation at oncogenic loci. To determine whether KAT2A itself organizes into nuclear condensates in AML cells, we performed immunofluorescence for KAT2A in AML and non-AML cell lines. KAT2A forms discrete nuclear puncta consistent with phase-separated condensates in all examined cell lines (Figure 5H and Figure S4). Expectedly, treatment of OCI-AML3 cells with GSK699, the active cis-(R,R) enantiomer of GSK983 led to dose-dependent dissolution of KAT2A puncta, but strikingly, also to a near-complete elimination of ENL condensates but not total ENL protein levels in all GSK699-treated cell lines (Figure 5H-I and Figure S4&S5), despite ENL not being a direct substrate of GSK699. Given that ENL is important for the maintenance of oncogenic transcriptional programs in AML ^25,26^, our finding that KAT2A degradation dismantles ENL condensate formation suggests that KAT2A activity is a prerequisite for this process, positioning KAT2A/B activity as an upstream determinant of ENL-dependent transcriptional amplification.

### KAT2A/B degradation leads to a reduction in acetylation of core SEC members

The acetyltransferases KAT2A and B are known to not only acetylate histone substrates but also several non-histone proteins including prominent oncoproteins ^27^. We therefore performed acetyl-proteomics using TMT-based quantification of acetylated peptides in MOLM-13 cells treated with GSK983 for 2 hours (Figure 6A). Unexpectedly, in this study, among the most significantly depleted acetylated peptides were those mapping to three core members of the Super Elongation Complex - AFF1, AFF3, and ENL – even as their total protein levels remained unchanged (Figure 6B-C and Table S8). Sites that specifically lost acetylation were ENL at lysine 252 (K252), AFF1 at lysine 610 (K610) and AFF3 at lysine 563 (K563). Given the profound effects of KAT2A/B degradation on chromatin occupancy of ENL as well as on transcription from ENL occupied gene loci, we examined whether KAT2A/B activity is required for ENL to maintain its biochemical interactions with SEC complex members and chromatin-associated partners. To address this, we performed FLAG immunoprecipitation (IP) from HEK293T cells stably expressing FLAG-ENL or a FLAG-ENL in which the lysine 252 site was mutated to an alanine (ENL K252A mutant) and subjected these to mass spectrometry analysis of co-purified proteins. We specifically assessed the nuclear fraction to enrich for chromatin-associated interactions. Our results demonstrated that FLAG-ENL pulldown robustly enriched several SEC components including AFF1, AFF3, AFF4 and ELL (Figure 6D), as well as the H3K79 methyltransferase DOT1L and the DOT1L complex (DotCom) members AF10/MLLT10 and AF17/MLLT6, and members of the PRC1 complex such as CBX1 as previously described ^28,29^ (Figure 6D). In addition, we identified several members of the DNA repair complex as ENL interactors, and also identified all three components of the casein kinase 2 (CK2) complex. Strikingly, compared to the wildtype ENL, there was a substantial reduction in enrichment of several components of the super elongation complex including AFF1, AFF3, AFF4 and ELL in the ENL K252A mutant IP, while other key interactions such as with DotCom remained largely unchanged (Fig. 6E). These results indicate that ENL acetylation at lysine 252 regulates SEC assembly., Taken together with our findings that KAT2 degradation leads to loss of ENL K252 acetylation, ENL puncta formation and ENL chromatin occupancy,

**Figure 6.**
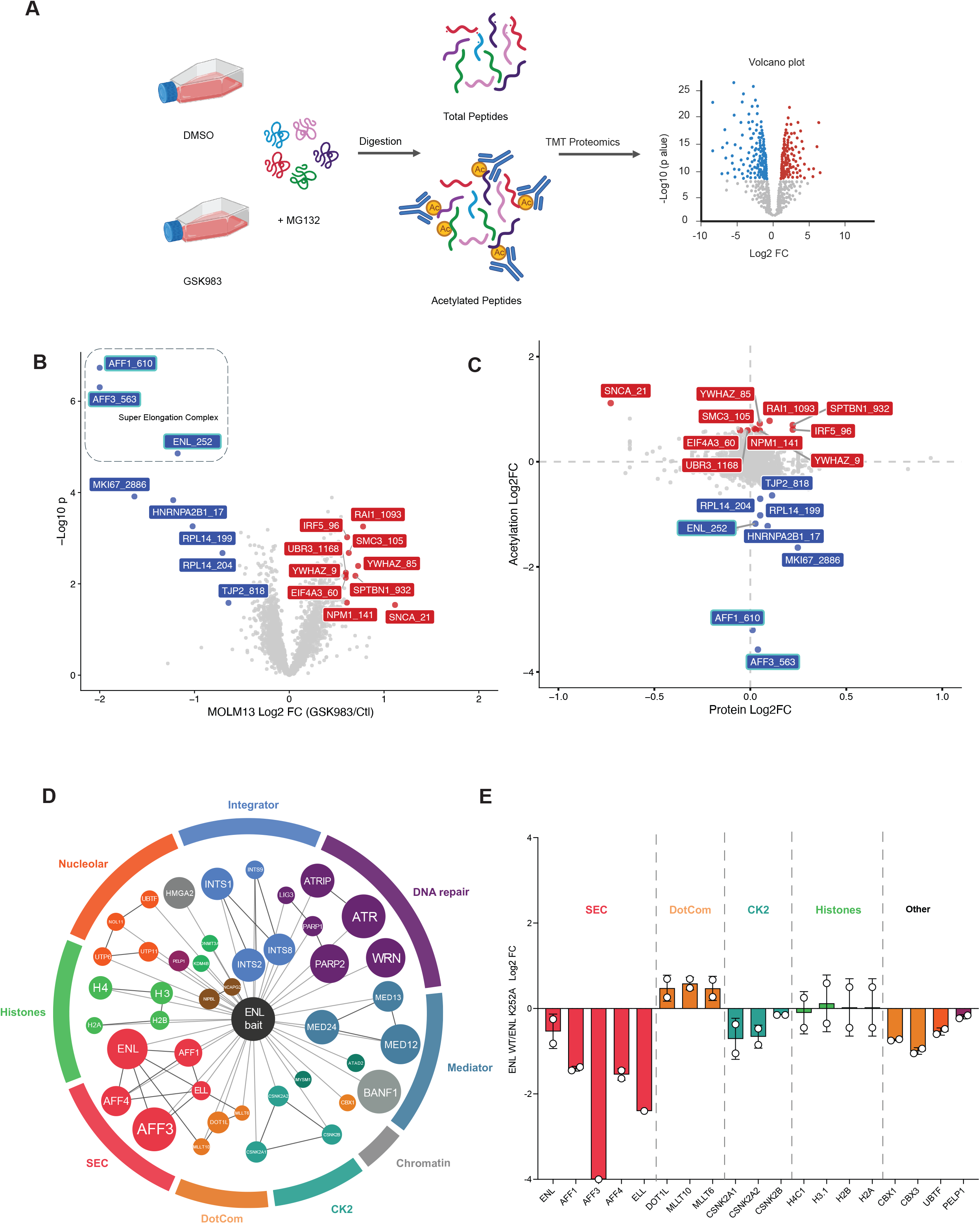
GSK983-mediated KAT2A/B degradation reduces SEC acetylation and disrupts ENL-SEC chromatin interaction. **(A)** Schematic of the quantitative acetyl-proteomics workflow. MOLM-13 cells were treated 700 with GSK983 or DMSO in the presence of MG132 to prevent proteasomal clearance of 701 acetylated peptides, followed by parallel TMT-based quantification of total and acetylated 702 peptides. A representative volcano plot of global protein abundance changes (GSK983 vs. 703 DMSO) is shown. **(B)** Volcano plot of acetylated peptide abundance in MOLM-13 cells treated with GSK983 705 relative to DMSO (x-axis, log_2_ fold change; y-axis, −log_10_ p-value). Peptides with decreased 706 acetylation including those corresponding to Super Elongation Complex (SEC) members AFF1, 707 AFF3, and ENL (MLLT1) are highlighted in blue whereas peptides with increased acetylation are highlighted in red. Dashed box denotes SEC-associated acetylated peptides. **(C)** Scatter plot comparing protein-level abundance changes (x-axis, protein log_2_FC) versus acetylation-site changes (y-axis, acetylation log_2_FC) for peptides identified in the acetyl-proteomics dataset. Red labels indicate peptides with increased acetylation; blue labels indicate 712 peptides with decreased acetylation and grey indicates unchanged acetylation. Residues with 713 changed acetylation are indicated with the amino acid code and the residue number (e.g. 714 ENL/MLLT1 at lysine 252 (ENL K252)). **(D)** Scatter plot of Flag-ENL co-immunoprecipitation mass spectrometry data from HEK293T 716 cells treated with GSK983 or DMSO. X-axis shows log_2_ enrichment of proteins in Flag-ENL 717 wildtype (WT) pulldown relative to Flag control (nuclear fraction); y-axis shows log_2_ fold change of FLAG-ENL K252A mutant IP relative to Flag-ENL WT pulldown. Bar groups are color-coded by protein complex/functional category as indicated in the complex name labels.

Our results establish that pharmacological degradation of KAT2A/B dismantles a hierarchical epigenomic dependency in AML - collapsing H3K9ac at identity-defining oncogene loci, displacing ENL-anchored transcriptional condensates from chromatin and suppressing MYC-driven proliferative and metabolic programs - with broad anti-leukemic activity that extends across genetically diverse AML subtypes regardless of the underlying oncogenic driver.

## DISCUSSION

The dysregulation of chromatin-modifying complexes has been recognized as a hallmark of malignancy, with the therapeutic targeting of epigenetic regulators emerging as a compelling strategy across multiple cancers ^30^. Apart from recurrently mutated chromatin regulators, it is now appreciated that malignant cells can acquire profound dependencies on chromatin complexes that are themselves not altered in cancer - so called non-oncogene dependencies - where the normal chromatin machinery is co-opted to sustain oncogenic transcriptional programs. Several groups including ours have shown that identifying such dependencies through unbiased, genome-scale approaches can reveal cancer-selective vulnerabilities in AML ^13,15,31^. Prior genetic studies including from our lab have identified several components of the SAGA complex including KAT2A and the chromatin reader protein SGF29 as a regulator of AML-associated transcription ^13–16,32^. In this study, through a comprehensive analysis of dependency data from > 1,000 cancer cell lines spanning diverse tumor types, we nominated the SAGA complex - a multi-subunit chromatin regulatory complex not recurrently mutated in AML - as one of the most significantly and selectively essential chromatin complexes in hematological malignancies. This selectivity, manifest across both myeloid and lymphoid lineages, defines SAGA as a non-oncogene dependency and provides a basis for its therapeutic targeting in hematological malignancies and particularly in AML.

Using a PROTAC-mediated strategy, we show that targeted degradation of KAT2A/B exerts potent, broad-spectrum anti-leukemic activity across genetically diverse AML cell lines, primary patient samples, and leukemia models – regardless of the underlying oncogenic driver. Mechanistically, we establish that KAT2A/B activity is required to maintain the H3K9ac chromatin state at a select set of asymmetrically acetylated AML identity genes, as well as acetylation of key components of the super elongation complex, and that loss of KAT2A/B triggers a cascade of downstream events: displacement of the ENL chromatin reader, dissolution of ENL-anchored transcriptional condensates at oncogenic loci, and coordinate suppression of MYC-driven proliferative and metabolic programs. Together, these findings delineate a hierarchical epigenomic circuit in which KAT2A/B functions as an upstream gatekeeper of leukemic transcriptional identity. Our CRISPR screens demonstrate that ENL is the key downstream effector of KAT2A/B activity in AML. ENL is a chromatin reader protein whose YEATS domain engages acylated histone marks - including H3K9ac - at active gene promoters, recruiting the super elongation complex to drive productive RNA Pol II elongation. Our data establish that KAT2A/B-deposited H3K9ac is required for ENL occupancy at AML oncogene promoters, and that KAT2A/B degradation results in near-complete dissolution of ENL nuclear condensates without affecting total ENL protein levels. This finding positions KAT2A/B as an upstream determinant of ENL-dependent transcriptional amplification in AML. Importantly, ENL loss itself confers resistance to GSK983, indicating that ENL displacement is not merely a correlate but a functional requirement for anti-leukemic effects of KAT2A/B depletion. The acetyl-proteomics data extend this mechanistic model by revealing that loss of KAT2A/B from AML cells leads to rapid and significant reduction of acetylation of the core members of the super elongation complex, including ENL at K252, AFF1 at K610, and AFF3 at K563. Loss of these acetylation marks occurs within one hour of GSK983 treatment, preceding any overt changes in SEC assembly, suggesting that regulation of SEC acetylation by KAT2A/B may cooperate with H3K9ac to stabilize ENL-chromatin or ENL-SEC interactions. The observation that compared to the wildtype ENL, the FLAG-ENL K252A mutant shows a significantly diminished interaction with multiple SEC components is consistent with the model thatKAT2A/B-dependent acetylation licenses the assembly of the super elongation complex.

Taken together, our study provides a comprehensive pharmacological and mechanistic evaluation of KAT2A/B degradation as a therapeutic strategy in AML. Despite dual KAT2A/B degradation, we establish degrader selectivity across several hundred cancer cell lines, characterize activity across multiple genetically distinct AML subtypes, primary patient samples, and patient-derived xenografts, and identify a strikingly consistent transcriptional signature of KAT2A/B loss shared across much of the genetic diversity of AML. Critically, we resolve the mechanistic basis for this activity in molecular detail, tracing a path from H3K9ac loss at AML oncogene loci to ENL eviction and dissolution of ENL-anchored transcriptional condensates at least in part through the acetyl-regulation of SEC complex assembly. This places KAT2A/B, and by extension the SAGA complex, at the interface of chromatin acetylation and transcriptional condensate biology. We propose that this SAGA-SEC connection may have broader implications, given the increasingly recognized role of acetylation-dependent SEC condensate formation in transcriptional control more broadly and also in SEC-dysregulated cancers ^25,33,34^. At the time of submission of this manuscript, a study reported a CRBN-based molecular glue with degron-independent, KAT2A-selective activity that similarly reduced H3K9ac and showed antileukemic effects^35^ reflecting a growing recognition of KAT2A/B as a druggable AML dependency. Our findings define the breadth of KAT2A/B dependency across AML genetics and resolve its downstream mechanistic consequences at the level of the SEC and ENL-anchored condensates in molecular detail.A key therapeutically significant aspect of our findings is the breadth of anti-leukemic activity demonstrated by GSK983 across AML subtypes that are otherwise mechanistically and genetically distinct. Current targeted therapies in AML - including menin inhibitors, FLT3 inhibitors, and IDH1/2 inhibitors — are largely restricted in their activity to defined molecular subgroups, and resistance commonly emerges through reactivation of the targeted pathway or engagement of alternative transcriptional programs. Our data demonstrate that KAT2A/B degradation retains full activity in AML subtypes which lack the HOX-driven transcriptional dependency (such as core binding factor AML), that limits the efficacy of menin and DOT1L inhibitors in this subtype. This mechanistic orthogonality reflects the fact that KAT2A/B-dependent acetyl regulation of histones and the SEC complex is a convergent epigenomic requirement across several AML subtypes, supporting the existence of a common epigenomic dependency that is not restricted to a few oncogenic contexts.

The pronounced activity of GSK983 against leukemic progenitors warrants particular attention. The near-complete suppression of colony-forming capacity in KMT2A-rearranged AML cells, even in the presence of cooperating oncogenic hits such as FLT3-ITD or NRAS-G12D that are independently associated with therapeutic resistance indicates that leukemic progenitor self-renewal might be particularly sensitive to KAT2A/B degradation. Strikingly, this suppression persists beyond the pharmacological half-life of GSK983 in the 12-day CFU assay, raising the possibility that even transient KAT2A/B degradation induces an irreversible epigenetic state change in leukemic progenitors. The depletion of HSC-, MPP-, and LMPP-like transcriptional populations in GSK983-treated primary AML cells, together with reductions in LSC17 and LSC47 stem cell scores, is consistent with this interpretation and supports a model in which KAT2A/B activity is required for the maintenance of the leukemic stem cell state.

In summary, these findings establish KAT2A/B degradation or inhibition as an attractive therapeutic strategy for the treatment of AML, providing a strong rationale for the clinical development of next-generation KAT2A/B inhibitors for evaluation in AML spanning diverse genetic subtypes, including high-risk and therapy-resistant disease.

## STAR Methods

### Histone Mass Spectrometry

OCI-AML3 cell line was treated with 500 nM GSK983 or DMSO, in 4 replicates for 48 hr. 50 million cells were harvested for histone post-translational modification (PTM) profiling by mass spectrometry. At 48 hr. cells were pelleted by centrifugation and nuclei were isolated using cold nuclear isolation buffer (NIB), as previously described by ^36^. Histones were acid-extracted from isolated nuclei using 0.2 M sulfuric acid (H_2_SO_4_) and subsequently precipitated with trichloroacetic acid (TCA). The resulting histone extracts were submitted for downstream mass spectrometry analysis.

### Proteomics

Human AML cell line, OCI-AML3 was treated with 150 nM GSK983 or DMSO in T75 mm tissue culture flasks in total of 60 mL medium up to 24 hrs. At 6 and 24 hr time points, 20 million cells from each replicate of both, DMSO and GSK983 treated group were spun down and washed three times with cold PBS. The pellets were then lysed with cold RIPA buffer containing protease inhibitor cocktail (Thermo Fisher Scientific) and universal nuclease (Fisher Scientific) at 1 in 1000 dilution on ice. The lysates were spun at 16,000 g at 4°C for 20’to remove cell debris. Supernatants containing proteins were transferred to new pre-chilled tubes for LC-MS/MS analysis.

### RNA-sequencing (RNA-seq)

Five human AML cell lines harboring different fusion proteins were treated with 500 nM of GSK983 or DMSO for 48 hr. Cells were plated at a density of 0.3 million per mL in a total of 5 mL medium in non-tissue culture treated 6-well plates. At 48 hr, cells were spun down and pelleted for RNA extraction. Total RNA was extracted using RNAeasy mini kit (Qiagen) as per the manufacturer’s protocol. 1 ug of total RNA was submitted for library prep followed by sequencing on NovaSeq X Plus series to obtain 150 bp pair end reads at Novogene.

### SLAM-seq

OCI-AML3 cells were treated with 150 nM GSK983 or DMSO, in 3 replicates for 4hr. to identify nascent RNA molecules being formed after KAT2A/B degradation by SLAM-seq. At 4 hr, cells were washed with cold PBS at 4°C for 5 min. Pellets were resuspended in pre-warmed RPMI medium containing 100 uM 4-thiouridine (S4U) and incubated in an incubator for 1 hr. Cells were then pelleted down for RNA isolation followed by alkylation using the SLAM-seq explorer kit (Lexogen) as per the manufacturer’s protocol. Library preparation for sequencing was performed using the QuantSeq 3’ mRNA-Seq V2 Library Prep Kit with UDI (Lexogen). The samples were sequenced on the Element Biosciences AVITI platform with the 2×75bp High Output Cloudbreak Freestyle Kit at SBP Genomics core, La Jolla.

### Drug screening analysis

Drug screening data analysis was performed in R (v4.5.1). For each time point and concentration, viability values of a replicate were flagged as outlier if it was ±2 SD from the mean of the group and set as missing value (NA). Outlier values were excluded from downstream calculations. For each sample, the mean viability of DMSO-treated replicates (after outlier removal) was used as the reference control (100% cell viability). The viability values of GSK983 treated replicates were normalized using the formula:

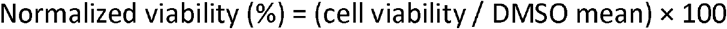

Two-sided Mann-Whitney U (Wilcoxon rank-sum) test for each concentration–sample–day combination. Multiple comparisons were corrected within each sample using the Benjamini-Hochberg method. Significance thresholds were: *** FDR < 0.001, ** FDR < 0.01, * FDR < 0.05. EC50 values were estimated using drc (v3.0.1) ^37^ by fitting a four-parameter log-logistic (4PL) dose-response model to mean normalized viability data using the drm() function, with the LL.4 model specification (parameters: Slope, Lower asymptote, Upper asymptote, EC50). EC50 estimates and 95% confidence intervals were extracted using the delta method .

### High Content Imaging

U2OS cells were seeded at a density of 1,500 cells/well, while OCI-AML3 and MOLM-13 cells were seeded at 10,000 cells/40 uL per well, in Poly-D-lysine-coated PhenoPlate™ 384-well microplates (Revvity Health Sciences) and allowed to adhere overnight. GSK699 was then dispensed in a dose-response range of 0.16 to 0.625 µM (1 in 2 dilutions) using an Echo 555 Acoustic Liquid Handler. Following incubation for 24 hours, cells were fixed with 4% paraformaldehyde (PFA) for 15 minutes and permeabilized with 0.2% Tween-20 in PBS for 30 minutes. After blocking with 3% BSA for 30 minutes, cells were incubated overnight with primary antibodies against KAT2A (Cell Signaling technologies) and ENL (Cell Signaling Technologies) at 1:200 and 1:100 dilutions respectively. The cells were then labeled with appropriate fluorophore-conjugated secondary antibodies (Invitrogen) at 1:2000 dilution and counterstained with Hoechst 33342 (Invitrogen) for 1 hour. Images were acquired using the Opera Phenix Plus High-Content System and analyzed with the Signals Image Artist software.

## ACKNOWLEDGEMENTS

We would like to thank Chih-Cheng Yang and Chun-Teng Huang from the Sanford Burnham Prebys Medical Discovery Institute (SBP) functional genomics core, the proteomics core, Yoav Altman from the SBP Flow Cytometry Core, and Rebecca Porritt and Kang Liu from the Genomics Core for their excellent support. We would also like to thank Dr. Akihiko Yokoyama for his helpful comments on the manuscript. We would also like to acknowledge the support of Wuxi Apptech (formerly HD Biosciences) for the BoE studies and Ryan Kunz and Brian Erickson from IQ Proteomics for their assistance with the acetyl proteomics experiments.

This work was supported by National Institutes of Health (NIH) National Cancer Institute grants CA262746, CA301721 and P30 CA030199, Rally and Luke Tatsu Foundation grant # 24IN24, the California Institute of Regenerative Medicine (CIRM) grant # DISC0-15816 and the American Cancer Society grant RSG-24-1320206-01-IBCD and the National Institutes of Health R50 CA211404 to M.W. We would also like to acknowledge the support of the Animal Facility, SBP Flow Cytometry Core, Genomics core, Functional Genomics core as well as the Bioinformatics core supported by the NCI Cancer Center Support Grant P30 CA030199.

## Author contributions

**AD** designed and performed experiments, analyzed and interpreted all data, prepared figures, and wrote the manuscript. **CYC, MP, NN and DF** designed and performed experiments, collected and processed experimental data, and contributed to data interpretation. **NS, EZ** and **RM** processed and analyzed data and provided computational support. **AS, MW** and **CJ** provided key reagents and resources critical to this study. **BAG** and **KV** provided critical conceptual input. **AJD** conceived and designed the study, supervised all experimental work, acquired funding, and wrote the manuscript with input from all authors. **AC, YY, KJP, AU, SO, DN & TP** contributed to the design and execution of acetyl proteomics experiment.

