## Supplemental Figures and Methods for "Disrupting a Convergent Acetylation Circuit Collapses Leukemic Identity Across AML Subtypes"

#### Figure 1

**A**

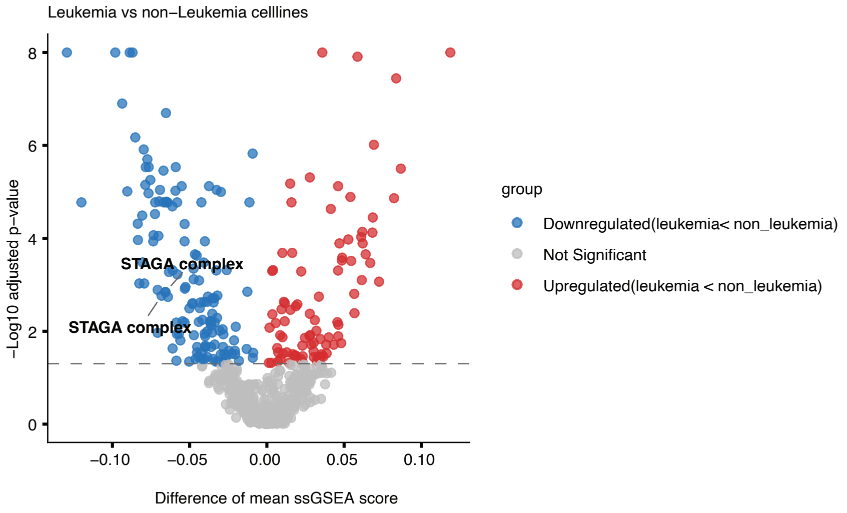


**B**

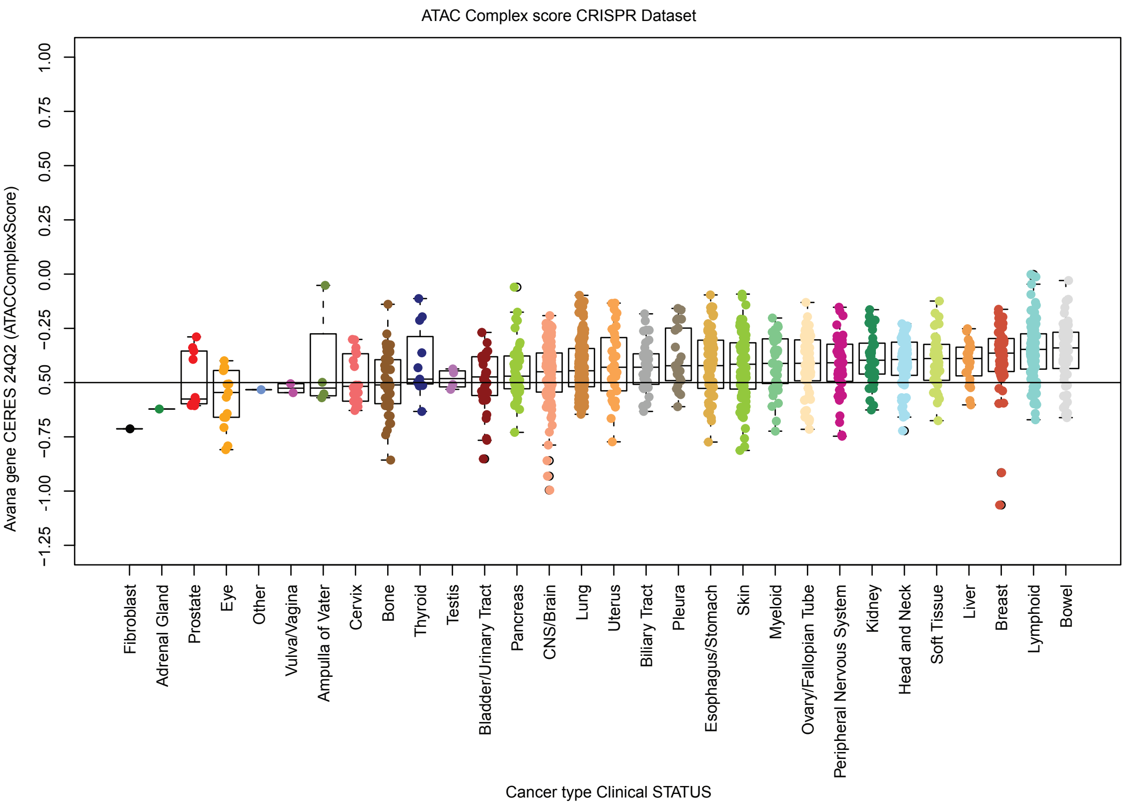

### Figure 2

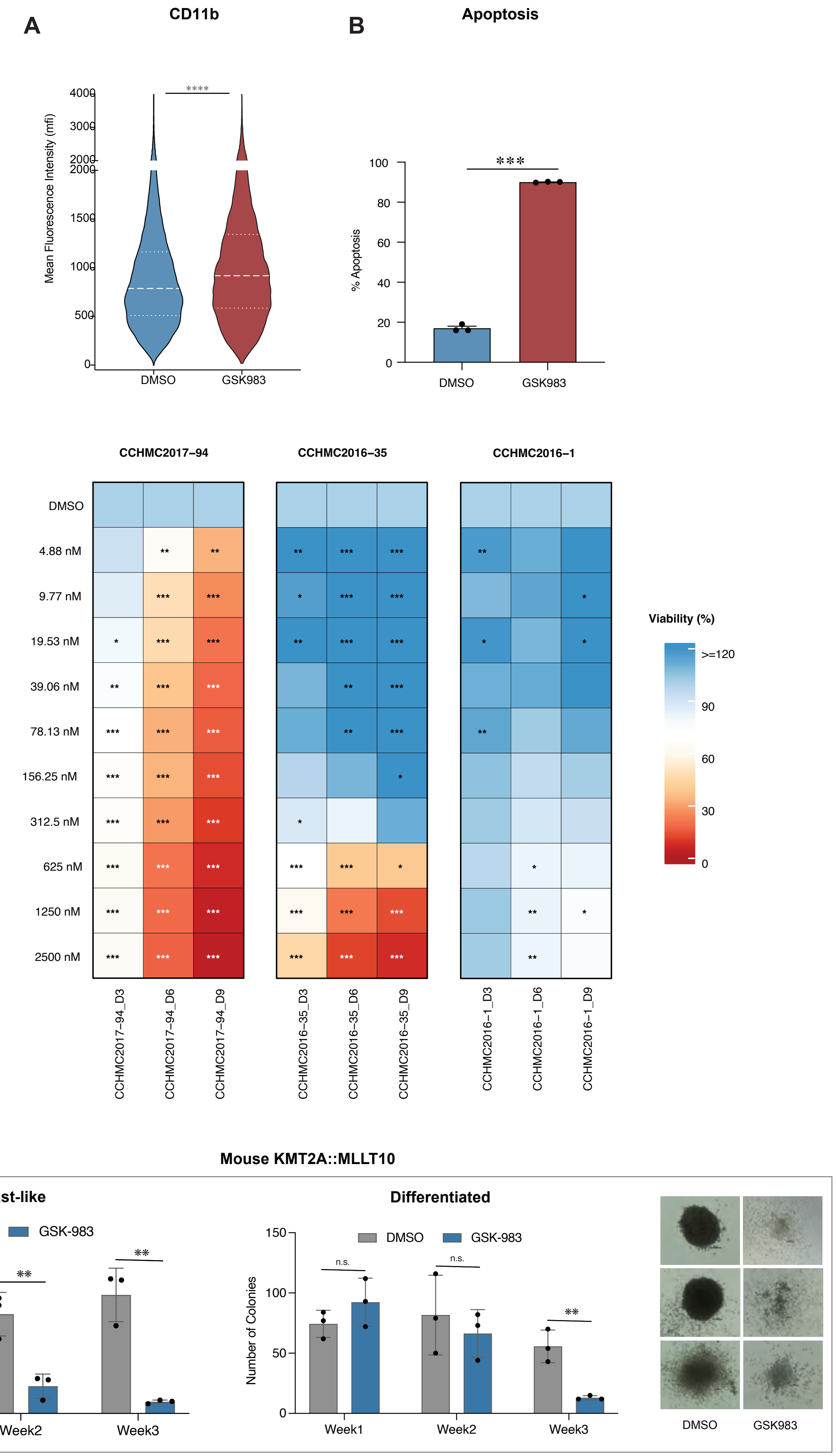
A
CD11b
Mean Fluorescence Intensity (mfi)
4000
3000
2000
1500
1000
500
0
****
DMSO
GSK983
B
Apoptosis
% Apoptosis
100
80
60
40
20
0
***
DMSO
GSK983
C
CCHMC2017–94
CCHMC2016–35
CCHMC2016–1
DMSO
4.88 nM
9.77 nM
19.53 nM
39.06 nM
78.13 nM
156.25 nM
312.5 nM
625 nM
1250 nM
2500 nM
Viability (%)
>=120
90
60
30
0
CCHMC2017–94_D3
CCHMC2017–94_D6
CCHMC2017–94_D9
CCHMC2016–35_D3
CCHMC2016–35_D6
CCHMC2016–35_D9
CCHMC2016–1_D3
CCHMC2016–1_D6
CCHMC2016–1_D9
D
Mouse KMT2A::MLLT10
Blast-like
Differentiated
DMSO
GSK-983
Number of Colonies
150
100
50
0
**
n.s.
Week1
Week2
Week3
DMSO
GSK983

Figure 3

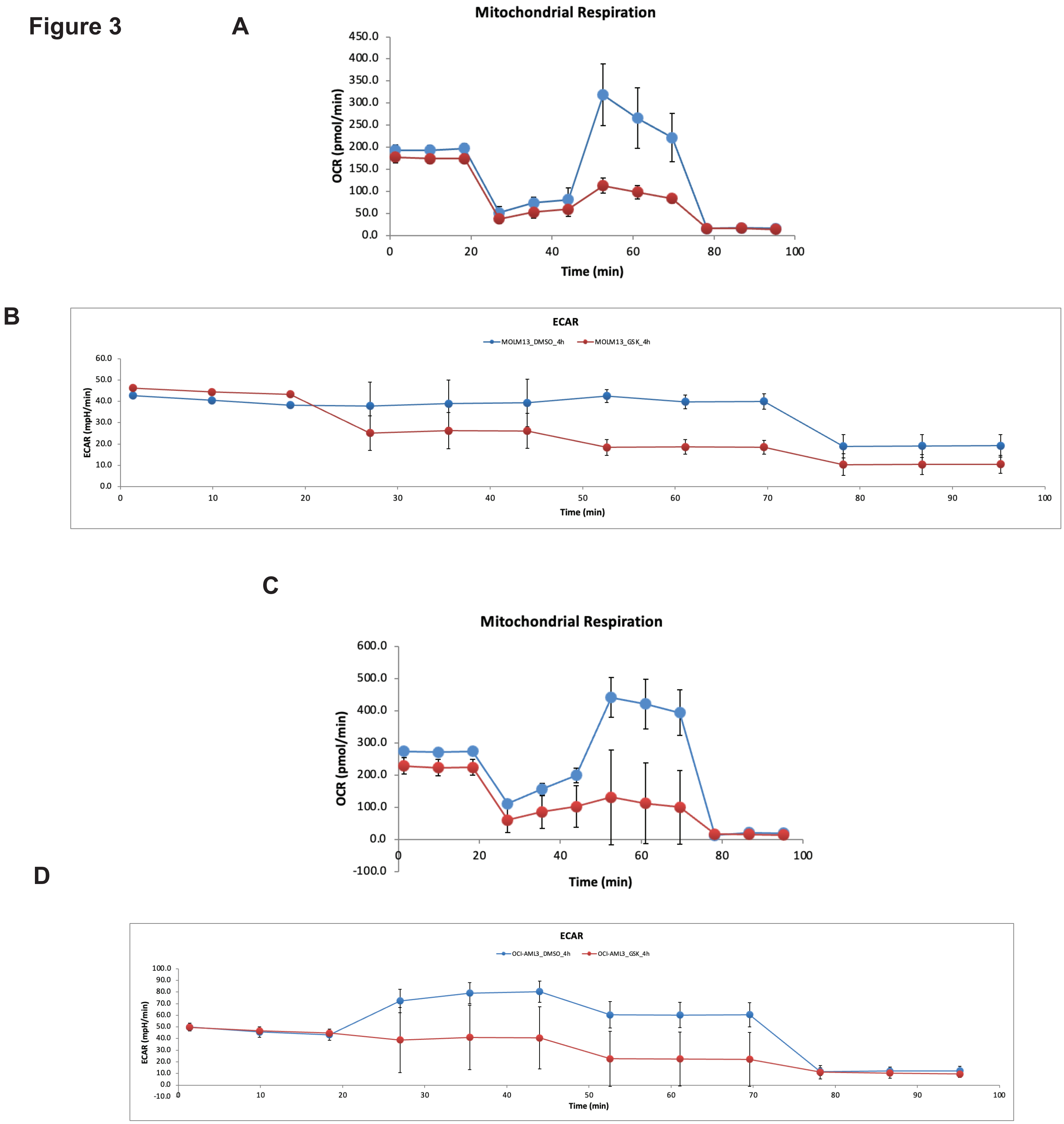
A
Mitochondrial Respiration
OCR (pmol/min)
Time (min)
B
ECAR
MOLM13_DMSO_4h
MOLM13_GSK_4h
ECAR (mpH/min)
Time (min)
C
Mitochondrial Respiration
OCR (pmol/min)
Time (min)
D
ECAR
OCI-AML3_DMSO_4h
OCI-AML3_GSK_4h
ECAR (mpH/min)
Time (min)

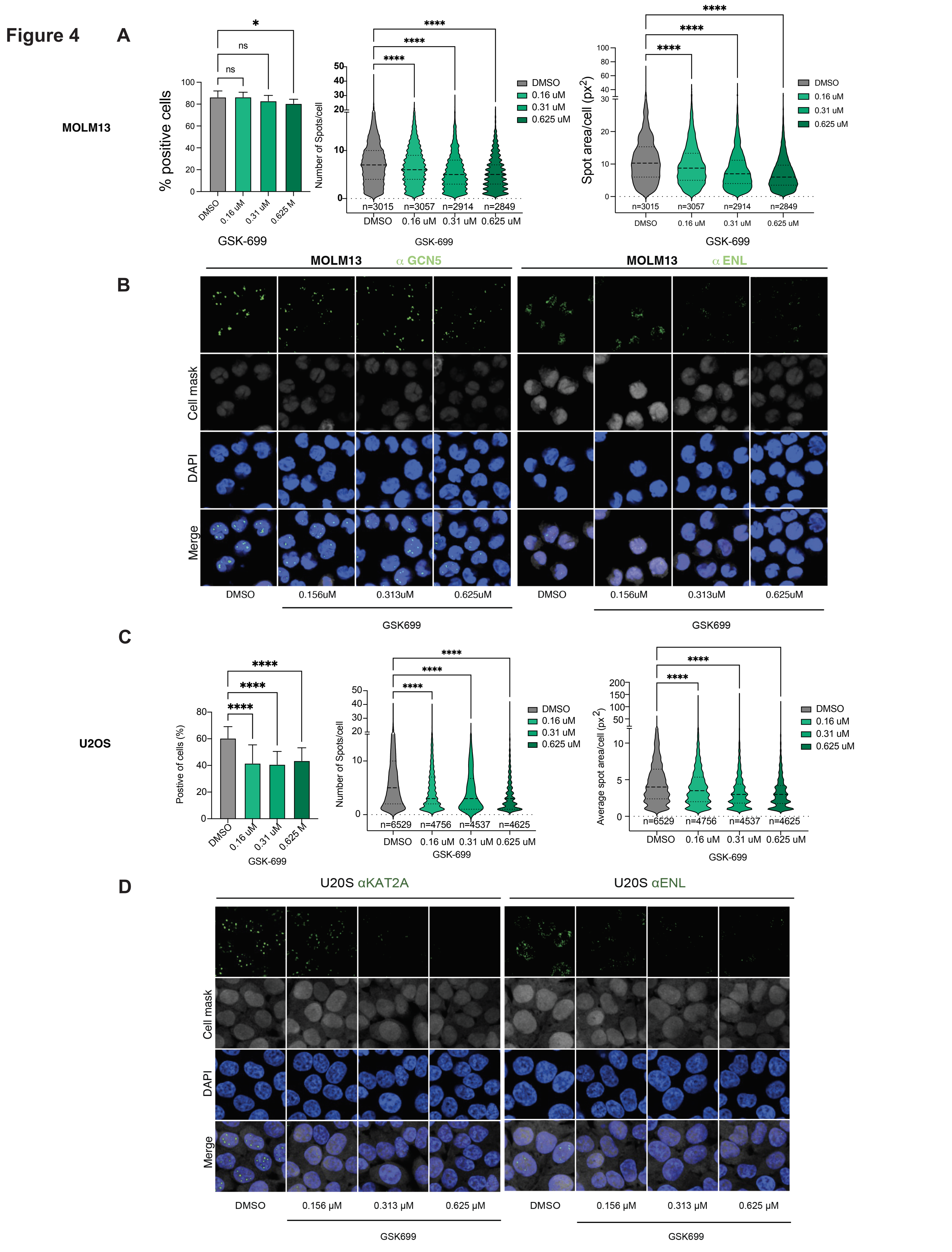
Figure 4
A
MOLM13
% positive cells
GSK-699
Number of Spots/cell
Spot area/cell (px2)
DMSO
0.16 uM
0.31 uM
0.625 uM
n=3015
n=3057
n=2914
n=2849
B
MOLM13 α GCN5
MOLM13 α ENL
Cell mask
DAPI
Merge
0.156uM
0.313uM
0.625uM
GSK699
C
U2OS
Postive of cells (%)
Average spot area/cell (px2)
n=6529
n=4756
n=4537
n=4625
D
U20S αKAT2A
U20S αENL
0.156 μM
0.313 μM
0.625 μM

**Figure 5**

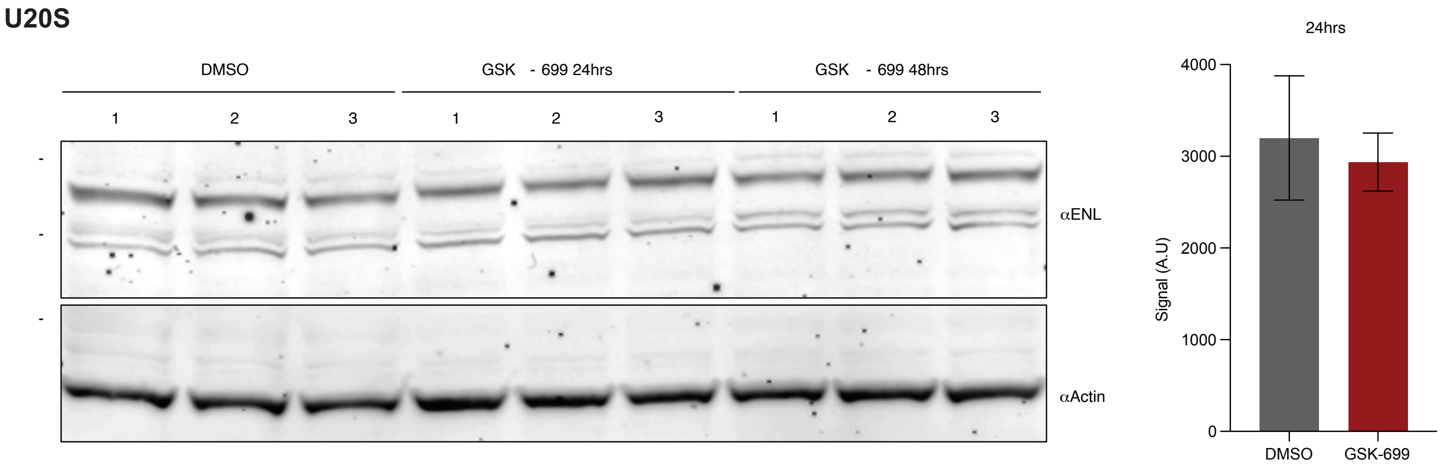
U20S
DMSO
GSK - 699 24hrs
GSK - 699 48hrs
1 2 3 1 2 3 1 2 3
αENL
αActin
24hrs
4000
3000
2000
1000
0
Signal (A.U)
DMSO
GSK-699

**Figure 1. SAGA but not ATAC  complex dependency is selective to leukemia.**

**(A)** Volcano plot of ssGSEA scores for 718 CORUM protein complexes comparing leukemia vs. non-leukemia DepMap cell lines. The SAGA complexes SAGA_475 and SAGA_476 from CORUM are labeled among the top leukemia-selective hits. Blue, leukemia-selective; red, selective in others; grey, non-significant. Dashed line indicates significance threshold.

**(B)** Boxplot of ATAC complex CERES dependency scores (Avana ATAC_Score) across cancer lineages. Each cancer type (names labeled below) are represented by unique colors. The solid grey line shows significance threshold.

**Figure 2. GSK983 promotes differentiation, apoptosis and impairs proliferation of genetically diverse AML models**

**(A)** CD11b surface marker expression (mean fluorescence intensity, MFI) changes in OCI-AML3 treated with DMSO or 5 uM GSK983 for 72 hr. Violin plots show distribution across 3 independent replicates with Mean fluorescence intensity of APC representing CD11b expression on the y axis. Thin and thick dashed lines indicate quartiles and median respectively.

**(B)** Apoptosis induction in OCI-AML3 treated with DMSO or 500 nM GSK983for 7 days. Percent apoptosis  quantified by flow cytometry is plotted on the y axis for DMSO and GSK983 treated cells Data are mean ± SEM; n = [3].

**(C)** Heatmap of GSK983 viability response across 3 primary AML patient-derived xenograft samples, CCHMC2017-94, CCHMC2016-35 (CBFA2T3::GLIS2) and CCHMC2016-1(PICALM::MLLT10) over the period of 9 days. Samples were treated with GSK983 at concentrations from 4.88 to 2500 nM shown in rows and viability is shown at Days 3, 6, and 9 as columns. Color scale for percent viability normalized to DMSO is shown on the right of the heatmaps. Statistical significance versus DMSO calculated by Mann-Whitney test is indicated as *p < 0.05, **p < 0.01, ***p < 0.001.

**(D)** Serial replating colony forming unit (CFU) assay over three consecutive weeks for mouse KMT2A::MLLT10 leukemia cells is shown in bar graphs with blast-line colonies on the left and differentiated colonies on right. Colonies per 500 cells were counted from 3 independent replicates every week and replated for counts on the following week. Grey bars represent colonies from DMSO treated whereas blue bars represent 900 nM GSK983 treated cells. Error bars represent SEM values; n.s.=non-significant, **≤0.05. Pictures of the representative colonies from DMSO (left panel) and GSK699 treated (right panel) are shown on the right. Scale bar;100 um.

**Figure S3. KAT2A/B degradation acutely impairs mitochondrial respiration and glycolysis in AML cells.**

(**A–B**) Seahorse XF Mito Stress Test in MOLM-13 cells treated with GSK699 or DMSO for 4 hours. Oxygen consumption rate (OCR; A) and extracellular acidification rate (ECAR; B) are shown over time (minutes). Sequential injections of oligomycin, FCCP, and rotenone/antimycin A are indicated.

(**C–D**) Corresponding OCR (C) and ECAR (D) measurements in OCI-AML3 cells under identical treatment conditions. Data are mean ± SEM. Blue and red lines represent DMSO control and GSK699 treatment respectively.

**Figure S4. Reduction of ENL condensate formation upon KAT2A/B degrader treatment.**

**(A) High-content immunofluorescence quantification of ENL nuclear condensates in MOLM13 cells treated with increasing doses of GSK699.** (Top, left) Percentage of ENL puncta-positive cells across different drug conditions (DMSO, 0.16, 0.31, and 0.625 µM GSK699) are shown. (Top, centre) The number of ENL puncta per cell is shown. (Top,right) The size of ENL puncta (spot area per cell, $px^2$) are shown as violin plots.*$p < 0.05$; ***$p < 0.001$

**(B)** Immunofluorescence (high content imaging) indicating changes in condensate formation of ENL after KAT2A degradation, GCN5/KAT2A (left, green) and ENL (right, green) in MOLM13 cells treated with DMSO, 0.156 µM, 0.313 µM, or 0.625 µM GSK699. Rows show results of staining with antibody against KAT2A and ENL (GFP, top row), Cell Mask (grey, second row), DAPI (blue, third row), and merged channels (bottom row). Representative images of the stained cells are shown.

**(C) High-content immunofluorescence quantification of ENL nuclear condensates in U2OS cells treated with increasing doses of GSK699.** (Top, left) Percentage of ENL puncta-positive cells across different drug conditions (DMSO, 0.16, 0.31, and 0.625 µM GSK699) are shown. (Top, centre) The number of ENL puncta per cell is shown. (Top,right) The size of ENL puncta (spot area per cell, $px^2$) are shown as violin plots.*$p < 0.05$; ***$p < 0.001$

**(D)** Immunofluorescence (high content imaging) indicating changes in condensate formation of ENL after KAT2A degradation, GCN5/KAT2A (left, green) and ENL (right, green) in U2OS cells treated with DMSO, 0.156 µM, 0.313 µM, or 0.625 µM GSK699. Rows show results of staining with antibody against KAT2A and ENL (GFP, top row), Cell Mask (grey, second row), DAPI (blue, third row), and merged channels (bottom row). Representative images of the stained cells are shown.

**Figure S5. Kat2A/B degrader treatment does not alter ENL protein expression.**

Immunoblot depicting ENL protein expression after treatment with GSK699 at 24 hr and 48 hr in U2OS cells is shown on right. Beta-actin was used as a loading control. N=3 independent experiments. Quantification of ENL protein bands at 24 hr using ImageJ is shown on the left.

#### SUPPLEMENTAL METHODS

**Cell culture:**

All human AML cell lines were cultured in RPMI 1640 medium supplemented with 2 mM l-glutamine and sodium pyruvate, 10% fetal bovine serum (FBS), and 50 U/mL penicillin/streptomycin (Thermo Fisher Scientific, Carlsbad, CA), and incubated in 5% $CO_2$ at 37°C. AML Patient Derived Xenograft (PDX) cells were cultured in IMDM with 20% BIT,175nM UM171 and 100ng/mL HsSCF, 10ng/mL HsIL3, 20ng/mL HsIL6, 20ng/mL HsGMCSF, 20ng/mL HsFLT3L and 20ng/mL Hs TPO and incubated in 5% $CO_2$ at 37°C. PDX cells from Cincinnati Children's Hospital Medical Center (CCHMC) were cultured in IMDM with 20% FBS and 10 ng/mL of the Human cytokines SCF, IL3, IL6, TPO and FLT3L. Murine leukemia cells were cultured in DMEM with 15% FBS, 2 mM l-glutamine, 50 U/mL penicillin/streptomycin and 20 ng/mL murine SCF, 10ng/mL murine IL6 and 10ng/mL murine IL3.

**Apoptosis assay:**

100,000 cells of each, MOLM-13 and OCI-AML3 were plated in 12 well non-tissue culture treated plates and treated with GSK983 at 500 nM concentration or DMSO for seven days. At day 7, cells were spun down and washed with cold PBS and stained with anti-Annexin antibody (BD Pharmingen, #550475) for 20 min on ice, away from light. The viability stain, Sytox Blue (Life Technologies, #34857) was added at 1:1000 dilution prior to analysis. Percent apoptosis was measured by flow cytometry using Fortessa X20 (BD Biosciences).

**Differentiation marker analysis:**

100,000 cells of each human AML cell line were plated in 12 well non-tissue culture treated plates and treated with GSK983 at 1 uM and 5 uM concentrations or DMSO for three days. On day 3, cells were spun down and stained with APC-anti CD11b antibody (eBiosciences # 17-0112-83) for 20 min. on ice. 1in 1000 dilution of Sytox Blue was added as a viability dye to each sample. Cells were analyzed by flow cytometry using Fortessa X20 (BD Biosciences).

**Cell growth assay:**

Cell growth assay in human AML cell lines and PDX cells was performed over the period of 12 days using CellTitre-Glo Luminescence assay. GSK983 compound was dispensed using Echo 555 liquid handler (Beckman Coulter) in varying concentrations. Cells were plated at 2000 cells/ 50 uL density in total of 50 uL volume in 384 well non-tissue culture treated plates at day 0. Equal volumes from each well were used for CellTiter-Glo assay as well as 25 uL volumes from the assay plate were replated with fresh medium and the compound every 3 days. Cell-Titer-Glo cell viability assay was performed as per the manufacturer's instructions. The assay was performed similarly for AML primary as well as PDX cells over the period of 9 days by plating 4000 cells/well on day 0. Percent DMSO was calculated for each dose of the compound at each time point.

**Colony Forming Unit (CFU) assay:**

For Colony formation assays of human CD34+ve cells in vitro transformed with KMT2A:MLLT3 with different mutational profiles (a kind gift from Dr. Wunderlich at CCHMC, Cincinnati, OH) 2000 cells were plated per 35mm dish in 3 replicates, in 1 mL H4100 base media (Stem Cell Technologies) with 20% FBS and human cytokines: SCF, TPO, FLT3, IL-3, IL-6, G-CSF, and GM-CSF (Peprotech) at 10ng/mL with DMSO or 500 nM GSK699. Colonies were counted on day 10. CD34+ve cells transformed with MLL-AF9 and RAS G12D mutant were plated similarly to other CD34+ve cell leukemias in H4100 medium without cytokines. PDX sample CCHMC2017-14 was thawed and cultured in IMDM medium supplemented with 20% FBS, 50 U/mL penicillin/streptomycin and human cytokines as mentioned earlier for 2 days. 400,000 cells were plated in 1 mL IMDM complete medium for pre-treatment with DMSO or 500 nM GSK699 3 days prior to CFU assay. On Day 3, cells were spun down and plated at 3000 cells/1 mL in H4100 medium with 20% FBS and human cytokines and DMSO or 500 nM GSK699 described earlier for CD34+ve transformed cells. Colonies were counted on day 10.

For murine leukemia CFU assays, KMT2A::MLLT10 transformed primary mouse leukemia cells were treated with DMSO or 900 nM GSK983 and plated at 500 cells per 35 mm dish, in triplicates, in 1 mL methocult M3234 medium supplemented with mouse cytokines, SCF, IL6 and IL3 at 20ng/mL, 10ng/mL and 6ng/mL concentration respectively. Colonies were counted every 7 days for up to 3 weeks. At week 1 and 2, after colony counts, cells from all three replicates were washed with PBS and pooled in one tube. Cells were counted from DMSO as well as PROTAC groups and 500 cells per dish were replated with freshly added DMSO or PROTAC.

**Chromatin Immunoprecipitation and sequencing (ChIP-seq):**

Human AML cell lines were treated in triplicates with GSK983 or DMSO for 48 hr. for studying changes in histone 3 acetylation at lysine 9 (H3K9ac) upon KAT2A/B degradation. ChIP-seq was performed as described earlier [1]. 1 million cells from each replicate were fixed using 1% formaldehyde for 5 min. at RT and sheared using Bioruptor as described earlier [1]. sheared chromatin was immunoprecipitated overnight with H3K9acetyl antibody (Abcam). The eluted DNA after reverse crosslinking was subjected to library preparation using NEBNext Ultra II DNA library prep kit for Illumina (New England Biolabs). Sequencing was performed on the Element Biosciences AVITI platform with the 2x75bp High Output Cloudbreak Freestyle Kit at SBP Genomics core, La Jolla.

**CUT&RUN:**

Changes in genome wide binding of ENL, RNApol II as well as RNApol II CTD ser2p upon treatment with GSK699 in human AML cell lines were assessed using CUTANA CHIC/CUT&RUN assay kit (Epicypher). AML cell lines were treated with 1 uM GSK699 or DMSO at 0.3 million per 1mL medium in total of 5 mL in non-tissue culture treated 6-well plates in duplicates for 48 hr. At 48 hr, cells from all the replicates were counted and 500,000 cells from each were mixed with 50,000 mouse leukemia cells (as internal standard) for each antibody. CUT&RUN assay was performed as per the manufacturer's protocol. Libraries for sequencing were prepared using NEBNext Ultra II DNA library prep kit for Illumina (New England Biolabs). Sequencing was performed on the Element Biosciences AVITI platform with the 2x75bp High Output Cloudbreak Freestyle Kit at SBP Genomics core, La Jolla.

**Single cell RNA-seq (scRNA-seq):**

AML PDX sample, ATACC10, harboring the KMT2A::MLLT4 fusion was cultured for 48 hours prior to treatment initiation. Cells ($3\times10^6$) were seeded in 4 mL of medium in non-tissue culture-treated 6-well plates and treated with 500 nM GSK983 or DMSO as control. After 72 hours, 3 million cells per condition were harvested and live cells were enriched using the Dead Cell Removal Kit (Miltenyi Biotec) per the manufacturer's protocol. Viable cells from both conditions were submitted for scRNA-seq using the GEM-X Universal 3′ Gene Expression v4 kit (10x Genomics) at the SBP La Jolla Genomics Core. Samples were sequenced on an Element AVITI.

##### Aggregate gene dependency scores of CORUM complexes using DepMap CRISPR data

DepMap CRISPR gene effect data and cell line information was downloaded from DepMap portal (25Q3) [2]. List of CORUM v5.1 [3] complexes were downloaded from the CORUM database website. Aggregate gene dependency scores for CORUM complexes with at least 5 genes were calculated ssGSEA method implemented in GSVA v2.4.4 [4] package in R v4.5.2.

##### SLAM-seq and bulk RNA-seq data processing and analysis

Bulk RNA-seq samples were processed using nf-core [5] rnaseq pipeline (v3.14.0) with "-profile singularity --aligner star_rsem --igenomes_ignore --genome null" parameters and Ensembl (v111) human reference genome GRCh38 (primary assembly) and gene annotations. Differential expression analysis was performed using nf-core differentialabundance pipeline (v3.14.0) with "--filtering_min_abundance 5 --filtering_min_proportion 0.50" parameters and Ensembl (v111) gene annotations.

SLAM-seq samples were processed using nf-core slamseq pipeline (v1.0.0) with "-profile singularity --read_length 85 --multimappers" parameters and Ensembl (v111) human reference genome GRCh38 (primary assembly), gene annotations, and 3′ UTR coordinates. The nf-core slamseq pipeline uses SlamDunk [6] for alignment, filtering, and counting of T-to-C converted reads. Converted (T-to-C) read counts for 3′ UTRs of each gene's isoforms were aggregated for downstream analysis. Differential expression analysis was performed in R version 4.5.2 using DESeq2 version 1.50.2 [7].

##### scRNA-seq data processing and analysis

Raw data was processed using Cellranger count (v9.0.1), human reference genome GRCh38.p14, and GENCODE (v45). GSK983 and DMSO samples had 21,885 and 22,355 mean number of reads per cell, respectively. Median number of genes detected per cell for GSK983 and DMSO samples were 4,373 and 4,524, respectively. Doublets were detected using Scrublet [8] provided as part of Demuxafy [9] singularity image. Downstream analysis of scRNA-seq was performed using Seurat (v5.4.0) [10] and R (v4.5.2). Seurat analysis was performed in v3/v4 mode (options(Seurat.object.assay.version = "v3")) for compatibility with reference mapping using bone marrow single cell RNA-seq map by Zeng et al [11]. Cellranger output count matrices for GSK983 and DMSO samples were converted to Seurat objects using CreateSeuratObject with "min.cells = 5, min.features = 500" parameters. Doublets and low-quality cells with mitochondrial content percentage >10% were removed. Seurat objects for both samples were merged. Data was normalized using NormalizeData. Cell cycle phases were determined using CellCycleScoring and S/G2M gene sets provided with Seurat.

**H3K9ac ChIP-seq analysis**

Paired-end reads were trimmed using Cutadapt (v5.0) with parameters "-j 12 -m 20 -O 4 -q 15 -a AGATCGGAAGAGCACACGTCTGAACTCCAGTCAC -A AGATCGGAAGAGCGTCGTGTAGGGAAAGAGTGT". Read quality was assessed using Fastqc (v0.12.1). Trimmed reads were first aligned against chrM of human genome version GRCh38.p14 primary assembly using Bowtie2 (v2.5.4) and parameters "--local -X 2000" to remove mitochondrial reads. Unaligned reads to chrM were mapped to human reference genome GRCh38.p14 primary assembly using Bowtie2 with parameters "--very-sensitive --no-discordant -X 2000". Read alignments were filtered to keep only high quality and properly paired reads using alignmentSieve from deepTools (v3.5.5) with parameters "--minMappingQuality 30 --samFlagInclude 2". Duplicate reads were identified and removed using Picard MarkDuplicates (v3.4.0) (http://broadinstitute.github.io/picard). Normalized read coverage tracks in bigwig format were generated using bamCoverage from deepTools with parameters "--binSize 10 --smoothLength 200 -p 12 --normalizeUsing RPGC --effectiveGenomeSize 2913022398".

**CUT&RUN analysis**

Human and mouse spike-in reads were classified and separated using Xenome (v1.0.0) [12]. During first stage, human and mouse combined genome index for Xenome classification was generated using Xenome's index command with "--kmer-size 25" parameter and human GRCh38 primary assembly (as graft) and mouse GRCm38 primary assembly (as host) genome fasta files. Human and mouse reads were separated using Xenome's classify stage. Mouse spike-in sequencing read percentage range was 12%-24%. Read trimming, alignment, and alignment filtering steps were the same as H3K9ac ChIP-seq methods described above. Normalized signal generation step using deepTools bamCoverage was also similar to H3K9ac with additional "--scaleFactor" parameter to scale each sample by normalization scale factors calculated using mouse spike-in read percentages.

**Global proteomics (TMT) data processing and analysis**

Global TMT proteomics data processing and analysis was performed using MSstatsTMT v2.16.0 [13] package in R v4.5.1. Spectromine processed data was converted to MSstatsTMT format using *SpectroMinetoMSstatsTMTFormat*. Peptide level quantifications were summarized to protein level quantifications using *proteinSummarization* function. Missing data imputation was performed as described previously [14]. Differential abundance comparisons were performed using *lmfit* and *eBayes* functions from Limma v3.64.3 [15] package.

**Pooled CRISPR screen:**

The pooled epigenetic CRISPR library designed by our group [1] was utilized for the pooled CRISPR screen for identifying interactors of GSK983. The library virus was produced in HEK 293T cells as described earlier [1]. 40 million MOLM13 cells stably expressing spCas9 were transduced with the library virus to achieve less than 30% transduction efficiency ensuring expression of single sgRNA per cell. 48 hr post transduction, puromycin was added to the cultures at 1 ug/mL concentration. At 72 hr puromycin selection, cells were spun down and resuspended in fresh medium without puromycin. After 48 hr., cells were counted and split into 2 flasks each containing at least 6 million cells to maintain 500x representation of each sgRNA based on the library size. DMSO was added to one and 150 nM GSK983 was added to the other flask for performing the screen. 6 million cells were pelleted down for genomic DNA isolation at T0. Cells from both culture flasks were counted and at least 6 million cells from each were replated in fresh medium at 0.3 million/mL density every other day to confirm 500x representation at any given point. On Day 14 of the screen (T14), cells from DMSO as well as GSK983 treatment group were counted and 12 million cells were pelleted down for T14 genomic DNA isolation. The screen was performed in three independent replicates. Genomic DNA from all the replicates were isolated using Zymopure DNA isolation kit and libraries were made for next-generation sequencing as previously described [1]. Sequencing was performed to obtain 150bp pair end reads using Novaseq 500 series by Novogene.

**Immunoprecipitation (IP):**

HEK293T cells were lentivirally transduced to stably express 3xFLAG-ENL or 3xFLAG empty vector control. Approximately 70 million cells were treated with either 1 uM of GSK699 or DMSO for 4 hours prior to harvesting. Cells were lysed in NETN100 buffer as described earlier by Ui et.al, 2015, Mol.Cell [16]. The lysates were sonicated at 20% amplitude (5s on/20s off for 5 cycles) and centrifuged at 1,000g at 4°C for 15 minutes to separate the soluble fraction. The pellet (insoluble fraction) was then resuspended in NETN300 buffer supplemented with 0.1% Pierce™ Universal Nuclease (Thermo Fisher Scientific) and incubated at 4°C for 1 hour. Following centrifugation at 13,000g for 15 minutes at 4°C, the resulting soluble fraction was immunoprecipitated with Apexbio Anti-FLAG Magnetic Beads, and the precipitated proteins were submitted for Mass Spectrometry at the Proteomics core at SBP, La Jolla.

**Methods for acetylomics:**

MOLM-13 cells were treated with DMSO or 100 nM GSK983 for 2 h under standard culture conditions, with four biological replicates per condition. Cell pellets were processed for TMTpro-based quantitative proteomics, including acetyl-lysine peptide enrichment and matched total proteome profiling. Fractionated peptides were analyzed by LC-MS3 on an Orbitrap Fusion Eclipse using SPS-MS3-based TMT quantification, and spectra were searched against a human UniProt database using SEQUEST with peptide and protein identifications controlled at 1% FDR.

Differential acetylation was assessed after proteome normalization using fold-change estimates and Welch t-tests, with replicate quality evaluated by PCA, relative standard deviation, and Pearson correlation analysis.

**Cell viability profiling:**

A panel of 473 cancer cell lines was screened in a two-dimensional proliferation assay to evaluate sensitivity to GSK983. Cells were treated with GSK983 using a 3-fold serial dilution series beginning at 30 µM and cultured for 7 days under standard assay conditions. Cell proliferation was quantified at endpoint using a CyQuant imaging-based readout. Dose–response relationships were generated for each cell line, and compound sensitivity was summarized by calculating $GI_{50}$ values and area under the curve (AUC).
